# Multi-scale simulations of the T cell receptor reveal its lipid interactions, dynamics and the arrangement of its cytoplasmic region

**DOI:** 10.1101/2021.01.11.425722

**Authors:** Dheeraj Prakaash, Graham P. Cook, Oreste Acuto, Antreas C. Kalli

## Abstract

The T cell antigen receptor (TCR-CD3) complex initiates T cell activation following recognition of peptides presented by Major Histocompatibility Complex (pMHC)-encoded proteins. The ligation of pMHC to TCRαβ induces Src family kinases activity via the cytoplasmic tails of the CD3δε, CD3γε and ζζ dimers. The TCR-CD3 topology is well understood, but little is known about its conformational dynamics and arrangement of its cytoplasmic tails, limiting our grasp of the signalling mechanism. Here, we investigated the entire TCR-CD3 embedded in an asymmetric lipid bilayer using molecular modelling and multi-scale molecular dynamics simulations. Our study demonstrates conformational changes in the extracellular and transmembrane domains, and the arrangement of the TCR-CD3 cytoplasmic tails. The TCRαβ variable regions were the most flexible in the extracellular domain. The cytoplasmic tails formed highly interlaced structures while some tyrosine sidechains within the immunoreceptor tyrosine-based activation motifs (ITAMs) of the CD3ε and ζ subunits dynamically penetrated the hydrophobic core of the bilayer. Ionic interactions between the cytoplasmic tails and phosphatidylinositol phosphates (PIP_2_ and PIP_3_) in the inner leaflet of the lipid bilayer led to the formation of a distinct annular lipid fingerprint around the TCR-CD3 complex. These results combined with available experiential data increase our understanding of the TCR-CD3 activation mechanism and highlight the importance of membrane lipids in regulating T cell activation.

**Significance statement:** The T cell receptor (TCR-CD3) detects antigenic peptides displayed by major histocompatibility complexes (pMHC) to instigate activation of T cell adaptive immunity. Despite significant structural and functional knowledge of TCR-CD3 topology, the membrane interactions and dynamics of its cytoplasmic moieties remain elusive. Interactions of TCR-CD3 cytoplasmic tails with membrane lipids may regulate their phosphorylation by Src-family kinases, the first intracellular event required for T cell activation. Using the static 3D structure of TCR-CD3 resolved by cryo-electron microscopy, we provide novel insights into the protein-lipid interactions of the complete TCR-CD3 embedded in a bilayer closely mimicking its native membrane environment. Our study sheds light on the dynamics of the TCR-CD3 at near-atomic resolution and further aids in deciphering its activation mechanism.

## Introduction

T lymphocytes express a diverse repertoire of antigen receptors known as the T cell receptor (TCR-CD3 complex) on their plasma membrane. TCR identifies peptide antigens displayed in the jaws of major histocompatibility complexes (MHC) (1, 2). The TCR-CD3 complex consists of four non-covalently assembled dimers: TCRαβ, CD3δε, CD3γε heterodimers and the ζζ homodimer (3). Disulphide bridges aid in linking the subunits within the αβ and ζ ζ dimers in the extracellular region. Inter-subunit interactions are mediated by the ectodomains (ECDs) and the transmembrane regions (TMRs) which contribute to the stability of the complex and determine its precise topology (3-5). The α and β subunits each display variable domains, Vα and Vβ, featuring three variable loops of complementarity-determining regions (CDRs) 1, 2 and 3 that together form the VαVβ binding site for peptide-MHC (pMHC) ligands (6). The cytoplasmic regions (CYRs) of the α and β subunits each contain short peptides of less than ten amino acids long and do not transmit signals to the intracellular region.

TCR-CD3 signal transduction, which is initiated by pMHC binding to TCRαβ, is governed by the phosphorylation of immunoreceptor tyrosine-based activation motifs (ITAMs) in the intracellular region of CD3 and ζ subunits (7). Circular dichroism (CD) spectroscopy experiments suggested that the CYRs of each ITAM-containing subunit are intrinsically disordered in both monomeric and oligomeric states, but exhibit lipid-binding with acidic phospholipid containing vesicles (8). CD3ε and ζ CYRs contain basic-rich stretches (BRS) that have been suggested to mediate robust ionic interactions with negatively charged head-groups of phosphatidylinositols (9, 10) and phosphatidylserine (11) in the inner leaflet of the membrane. Moreover, the tyrosine sidechains in the ITAM-containing segments of both CD3ε and ζ CYRs were found to penetrate into the hydrophobic core of the membrane (11, 12). However, it remains unclear whether this configuration applies to all cytoplasmic tyrosines in an entire TCR-CD3 complex. A ‘stand-by’ model of TCR-CD3 signalling was proposed based on the evidence that a pool of constitutively active LCK at the T cell plasma membrane phosphorylates the ITAMs of TCR-CD3 CYRs upon their disengagement from the membrane (13). However, the molecular mechanism of pMHC binding that induces a change in the ITAM configuration which favours accessibility by LCK remains unclear. The key to these questions potentially lies in the interactions of the TCR-CD3 complex with its local membrane environment which is also currently poorly understood.

A recent cryo-electron microscopy (cryo-EM) study (4) revealed the 3D structure of the human TCR-CD3 complex (PDB:6JXR) at a resolution of 3.7Å and supports previous findings that TCRαβ maintains critical ionic contacts with CD3δε, CD3γε and ζ ζ in the TMR (3). This study shed light on the quaternary structure arrangement featuring highly interlaced contacts among subunits’ ECDs and TMRs, suggesting a dense connectivity maintaining the topology of the entire complex. However, the cryo-EM structure could not identify the arrangement of the CYRs presumably due to their disordered state. Moreover, the dynamic behaviour of the TCR-CD3 ECDs, TMRs and CYRs when embedded in their native membrane environment has not been studied. This information may provide clues to the signal transduction mechanism.

Here, we have used the cryo-EM structure to generate the first molecular model of the entire TCR-CD3 embedded in a complex asymmetric bilayer containing the predominant lipids found natively in its activation membrane domain (14). Our multi-scale molecular dynamics (MD) simulation approach, in coarse-grained and atomistic resolutions, provided insights into the conformational flexibility of the TCR-CD3 and its interactions with membrane lipids in the microseconds time-scale. We show that the CYRs assemble into a coiled conformation and interact with the inner membrane leaflet, while anionic head-groups of phoshadylinositol phosphate (PIP) lipids interact selectively with TCR-CD3 CYR. Our data also reveal that the ECDs, TMRs, and CYRs each exhibit conformational changes when the TCR-CD3 is embedded within the lipid bilayer, potentially supporting models of allosteric activation of the TCR-CD3 complex.

## Results

### Modelling the entire TCR-CD3 complex

We used the cryo-EM structure of the human TCR-CD3 complex (PDB:6JXR) as a template and performed secondary structure predictions with the PSIPRED 4.0 workbench (15) to model the entire complex. The predictions suggested that the cytoplasmic region (CYR) of the CD3δ, γ, ε subunits lacked secondary structure in agreement with circular dichroism experiments (8). However, the ζ CYR was predicted to contain short α-helices consistent with NMR spectroscopy data (16). These regions were modelled as helical in our complete TCR-CD3 structure while the rest of the ζ CYR were modelled as unstructured regions. Some extracellular residues in the CD3ε, CD3γ, TCRα subunits that were missing from the cryo-EM structure were also modelled as unstructured regions (**Suppl Fig S1**). The CYRs of the CD3 and ζ subunits were modelled in a linearly extended orientation perpendicular to the membrane to avoid bias in inter-subunit contacts at the beginning of the simulations. During our modelling, the TMRs and the ECDs of the TCR-CD3 subunits were position-restrained to preserve their experimentally derived structural integrity. Finally, twenty different models including each of their discrete optimized protein energy (DOPE) scores (17) were obtained. The structural models showed minimal differences from one another due to position restrains applied on the tertiary structure of the ECDs and TMRs, and on the predicted α-helices in both the ζ CYRs. The most energetically favourable model (least DOPE score) of the entire TCR-CD3 complex was used and further energy-minimised. This atomistic model (**Fig 1**) was converted to a coarse-grained (CG) resolution using the Martini forcefield and used for CGMD simulations.

**Figure 1.**
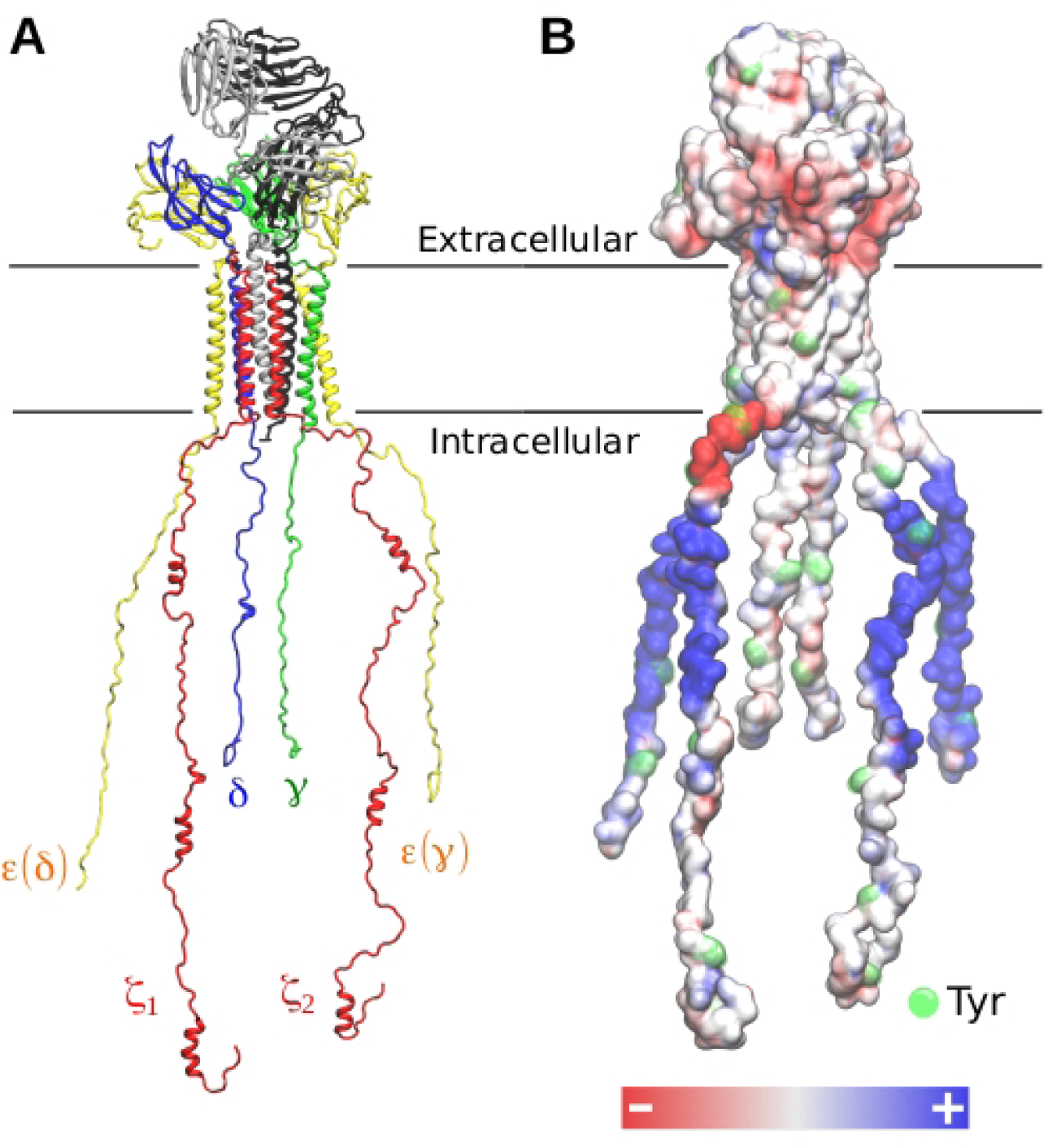
**(A)** Model of the entire T cell receptor used in our simulations. **(B)** Electrostatic profile of the TCR-CD3. Electronegative and electropositive regions are shown in red and blue respectively.

### The TCR-CD3 cytoplasmic region exhibits a coiled conformation

The CGMD simulations of the entire TCR-CD3 complex were used to analyse its dynamic nature when embedded in a lipid bilayer composed of the predominant lipids found in the TCR-CD3 activation domain (**Table 1**) (14). Five independent CGMD simulations of the TCR-CD3 were performed in an asymmetric complex membrane for 5 µs each. During the simulations, the CYRs of ζ ζ and CD3 dimers that were initially modelled in an extended configuration, rapidly coiled forming inter-chain interactions and then associated with membrane lipids of the inner leaflet (**Fig 2A**). Calculation of the distance between the center of mass (COM) of the CYRs and the COM of the lipid bilayer as a function of time suggested that the association of the CYRs with the membrane occurred within 1 µs of the simulations (**Fig 2B**). Although the time taken for the CYRs to coil and associate with the membrane was consistent amongst all CGMD simulations, their radius of gyration varied indicating that the coiled conformation of the CYRs is dynamic (**Fig 2C**). Due to the fluctuation in the CYR assembly, we wanted to determine its most commonly occurring structural conformation. Therefore, using the coiled and membrane-bound state of the CYRs (from the last 4 µs), we extracted 10,000 cytoplasmic configurations from all simulations combined and grouped them into clusters using a 3.5Å RMSD cut-off (RMSD calculated relative to our modelled structure shown in **Fig 1**). The largest cluster contained 867 similar structures (**Suppl Fig S2**) representing the most frequent structural conformation of the CYRs across all simulations (shown in a box in **Fig 2A**).

**Table 1.** Composition of lipids (%) in each leaflet of the membrane in all simulations.

| Lipid concentration (%) | POPC | POPE | SM | CHOL | POPS | PIP<sub>2</sub> | PIP<sub>3</sub> |
| --- | --- | --- | --- | --- | --- | --- | --- |
| Outer leaflet | 50 | 10 | 20 | 20 | - | - | - |
| Inner leaflet | 10 | 40 | - | 20 | 20 | 8 | 2 |

**Figure 2.**
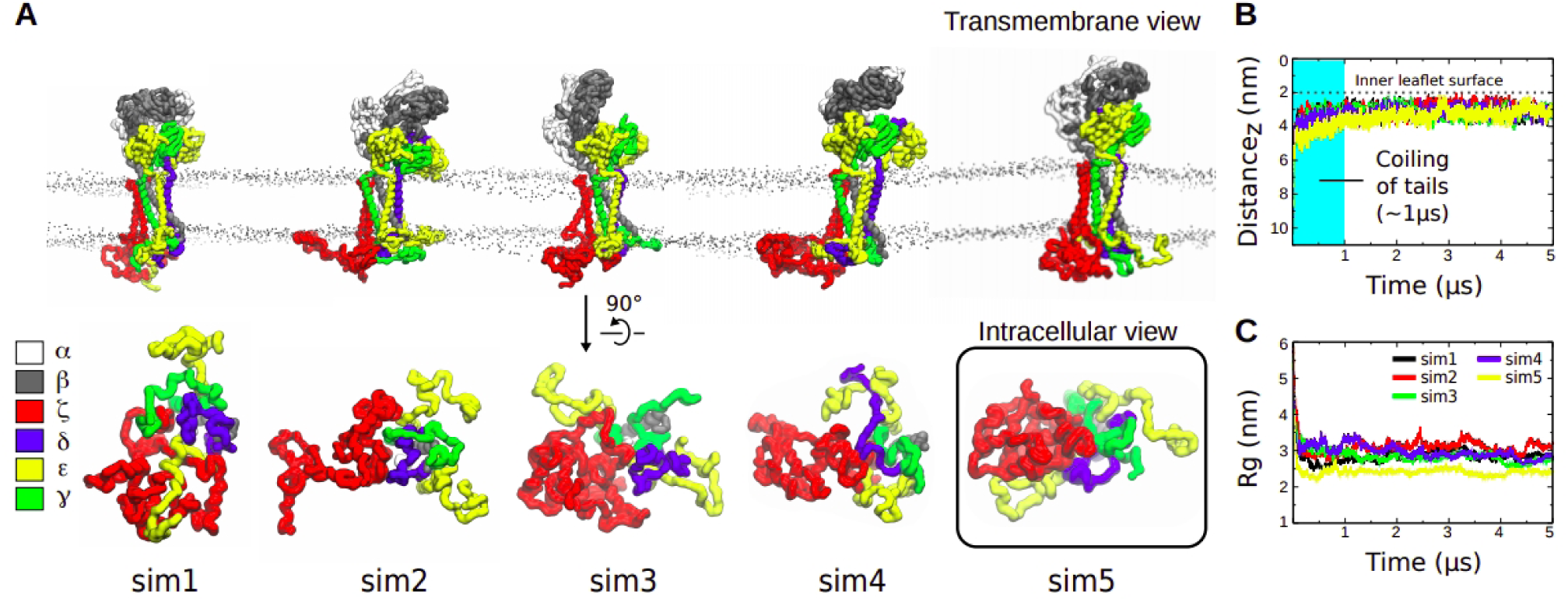
**(A)** Transmembrane view of the entire TCR-CD3 (top) and intracellular view of its cytoplasmic region (bottom) taken from the end of the five CGMD simulations. **(B)** Distance between the center of mass (COM) of the TCR cytoplasmic region and the COM of the membrane calculated from the five CGMD simulations of the entire TCR-CD3 versus time. **(C)** Radius of gyration (Rg) of the cytoplasmic region versus time. In (B) and (C), each coloured line represents one simulation.

### Membrane penetration by ITAM tyrosines and intracellular residues

Previous NMR studies of the CD3ε ITAM-containing peptide interacting with a lipid micelle suggested that two ITAM tyrosines, one isoleucine and one leucine residue penetrated the hydrophobic core of the membrane (11). To investigate whether our simulations showed similar membrane-penetrating activity of the ITAM tyrosines, we calculated their interactions with the hydrophobic acyl chains of lipids in our CGMD simulations. We found that membrane penetration was only achieved by some ITAM tyrosines. In all simulations combined, the tyrosines that displayed membrane-penetrating capabilities belonged to the CD3ε and ζ subunits only (**Fig 3A**). Y177 of CD3ε pairing with γ, referred to as CD3ε(γ), made the highest number of contacts with the lipid acyl chains (**Suppl Movie M1**), followed by Y177 of CD3ε(δ). Y166 of CD3ε(γ) penetrated the membrane more than Y166 of CD3ε(δ). In addition, only one of the subunits of the ζζ dimer mostly showed ITAM tyrosine contacts with lipid acyl chains. We also calculated the contacts of all TCR-CD3 subunits with the lipid acyl chains (**Suppl Fig S3A**). Interestingly, the short extracellular segments of both ζ subunits contacted the acyl chains of lipids in the outer membrane leaflet even more than their ITAM sequences, suggesting their tendency to anchor onto the extracellular leaflet during the simulations.

**Figure 3.**
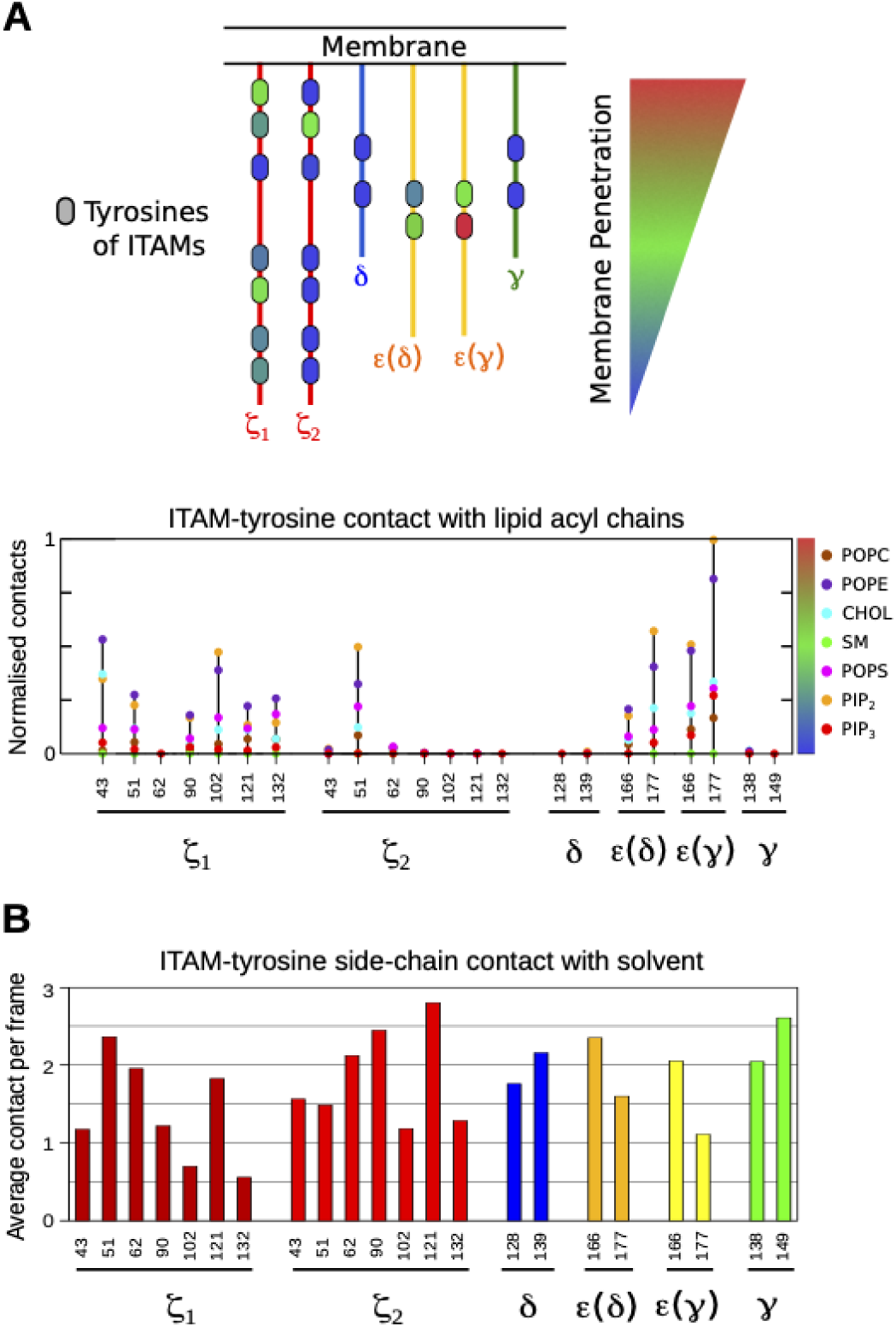
**(A)** Normalised number of contacts between the tyrosines of all cytoplasmic tails and the lipid acyl chains. Normalisation of contacts (N) was done by dividing all values (n) by the highest number of contacts (max_n) i.e. N=n/max_n. **(B)** Average number of contacts of each ITAM tyrosine with the solvent across 20 µs simulation time from all five simulations concatenated (1-4 µs each).

In order to study the solvent accessibility of ITAM tyrosines, we concatenated the last 4 µs of each of the five CGMD simulations when the CYRs were membrane-bound and calculated the average number of tyrosine sidechain contacts with intracellular water and ions (**Fig 3B**). We found that the ITAM tyrosines such as Y177 of CD3ε (γ) which penetrated the membrane the most interacted less with solvent as expected. However, ITAM tyrosines that did not show membrane penetrating ability also interacted less with the solvent suggesting that they dynamically shifted between hidden and exposed states within the coiled conformation of the CYRs. Nevertheless, the total solvent accessible surface area (SASA) of ITAM tyrosines in each ζ subunit was greater than that of the CD3δ, ε, γ subunits (**Suppl Fig S3B**).

### TCR-CD3 selectively interacts with lipid head-groups

We questioned if the association of the CYRs onto the inner leaflet of the bilayer affected the TCR-CD3 lipid environment. From all CGMD simulations combined, we analysed the contacts of the entire TCR-CD3 complex with all lipid head-groups including the sterol in both leaflets of the membrane, and further normalised the number of interactions of each lipid-type by their respective concentrations in the membrane. These data showed the relative enrichment of certain lipids over others in the vicinity of the TCR-CD3 complex. The TCR-CD3 showed a high propensity to contact phosphatidylinositol 4,5-biphosphate (PIP_2_) and phosphatidylinositol 3,4,5-triphosphate (PIP_3_) (**Fig 4A, 4B**) despite their relatively low abundance in the inner leaflet (8% and 2% respectively). A closer inspection revealed that the cationic residues dispersed throughout the CYRs of CD3ε and ζ subunits dominated the interaction with PIP_2_ and PIP_3_ lipids. The CD3δ and CD3γ CYRs also contacted the PIP lipid head-groups, though to a lower extent. In the CD3ε and ζ subunits, the basic residue-rich stretches (BRS) interacted most with PIPs, while the poly-proline motifs (PPPVPNP) in both CD3ε subunits (labelled ‘PPP’ in **Fig 4A**) showed the least contact. Basic residues at the TMR-CYR interface of the ζ subunits (R31, K33, R36) also contacted PIP lipids. Other anionic lipid head-groups i.e. POPS, and also neutral lipids made contacts with the CD3 and ζ CYRs but less frequently when compared with PIP lipids (**Fig 4A, 4B**).

**Figure 4.**
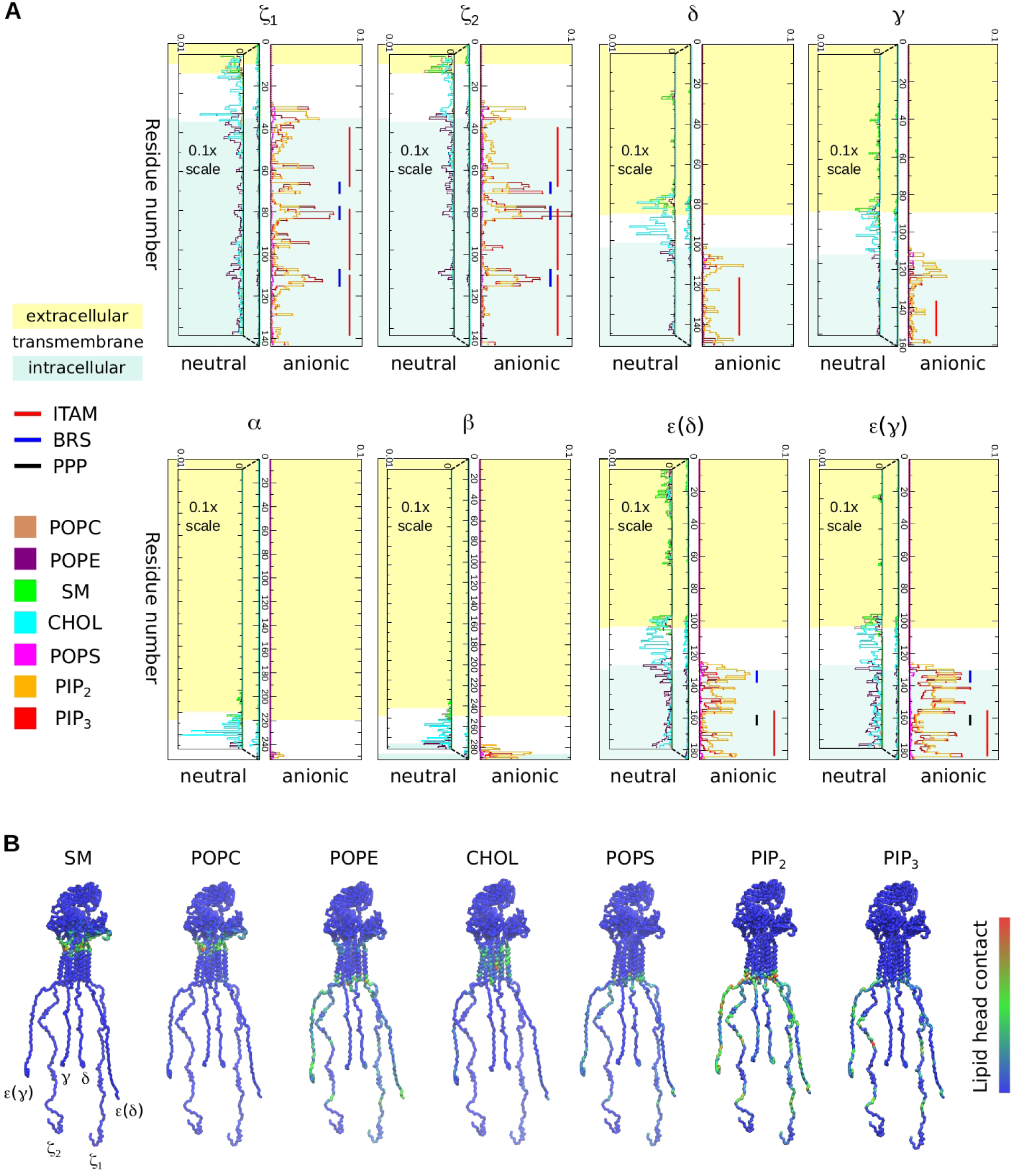
**(A)** Normalised number of contacts between the full-length TCR-CD3 subunits and lipid head-groups. The normalisation (N) is done by dividing the number of protein-lipid contacts of each residue (n1) by the number of the specific lipids in the bilayer (n2) and by the number of simulation frames (n3) i.e. N=n1/n2/ n3. Interactions of each TCR-CD3 subunit with anionic lipids are shown on the right whereas those with neutral lipids are shown on the left. The scale of non-anionic interactions was magnified 10 times (scale: 0 to 0.1) for clarity. ITAMs (red lines) on each subunit, BRS (blue lines) and poly-Proline motifs (black lines) of the CD3ε subunits are also indicated. **(B)** Contacts of the TCR-CD3 with lipid head-groups are mapped as a colour gradient (blue: low, green: medium, red: high) on the TCR-CD3 structure. The normalisation of contacts was done by dividing the contacts of each residue with a lipid type by the highest number of contacts with that lipid type.

Cholesterol interactions occurred across all the TMRs of the TCR-CD3, and were accompanied by minor interactions with CYRs of CD3ε and ζ (**Fig 4A**), explained by their ability to penetrate the membrane’s surface. POPE head-groups in the inner membrane leaflet also interacted with the CYRs of CD3ε and ζ to an extent similar to cholesterol interactions (**Fig 4A, 4B**). Sphingomyelin (SM), present only in the outer leaflet, interacted with all subunits but mostly contacted the extracellular segment of ζ and the ECD of CD3ε (δ), followed by the other CD3 subunits. We also observed that POPC head-groups in the outer leaflet interacted with the extracellular segment of ζ and with the connecting peptides (CPs) of CD3 subunits (**Fig 4A, 4B**). While in agreement with previous studies suggesting ionic interactions of CYRs with the plasma membrane (18), our data points to a potentially more complex scenario of contacts of the TCR-CD3 with membrane lipids than previously suggested.

### PIP clustering and the significance of the CYRs

Our investigation of PIP interactions with TCR-CD3 CYRs suggested clustering of PIPs around the TCR-CD3 complex. This led us to analyse the densities of each lipid type around the TCR-CD3. In the outer leaflet, there was no clustering of POPE, POPC, and SM, but in the inner leaflet the densities of PIP lipids around the protein dominated that of other lipids. To better discern the origin of this effect, we quantified the contribution of the ECDs, TMRs and CYRs of the TCR-CD3 toward lipid interaction. In addition to the CGMD simulations of the full TCR-CD3 complex (referred herein as FL), we performed two sets of CGMD simulations (5 simulations x 5 µs each) using the same membrane composition: (1) we excluded the ECDs and CYRs, and retained only the TMRs (referred herein as TMO), (2) we excluded only the CYRs, and retained the ECDs and TMRs (referred herein as ECTM) (**Fig 5A, Suppl Fig S4A**). We calculated the densities of lipids combining all five repeats of the TMO and ECTM simulations and compared them to the densities retrieved from FL simulations (**Suppl Fig S4B**). The most striking observation in all three conditions was the clustering of PIP lipids (**Fig 5B**).

**Figure 5.**
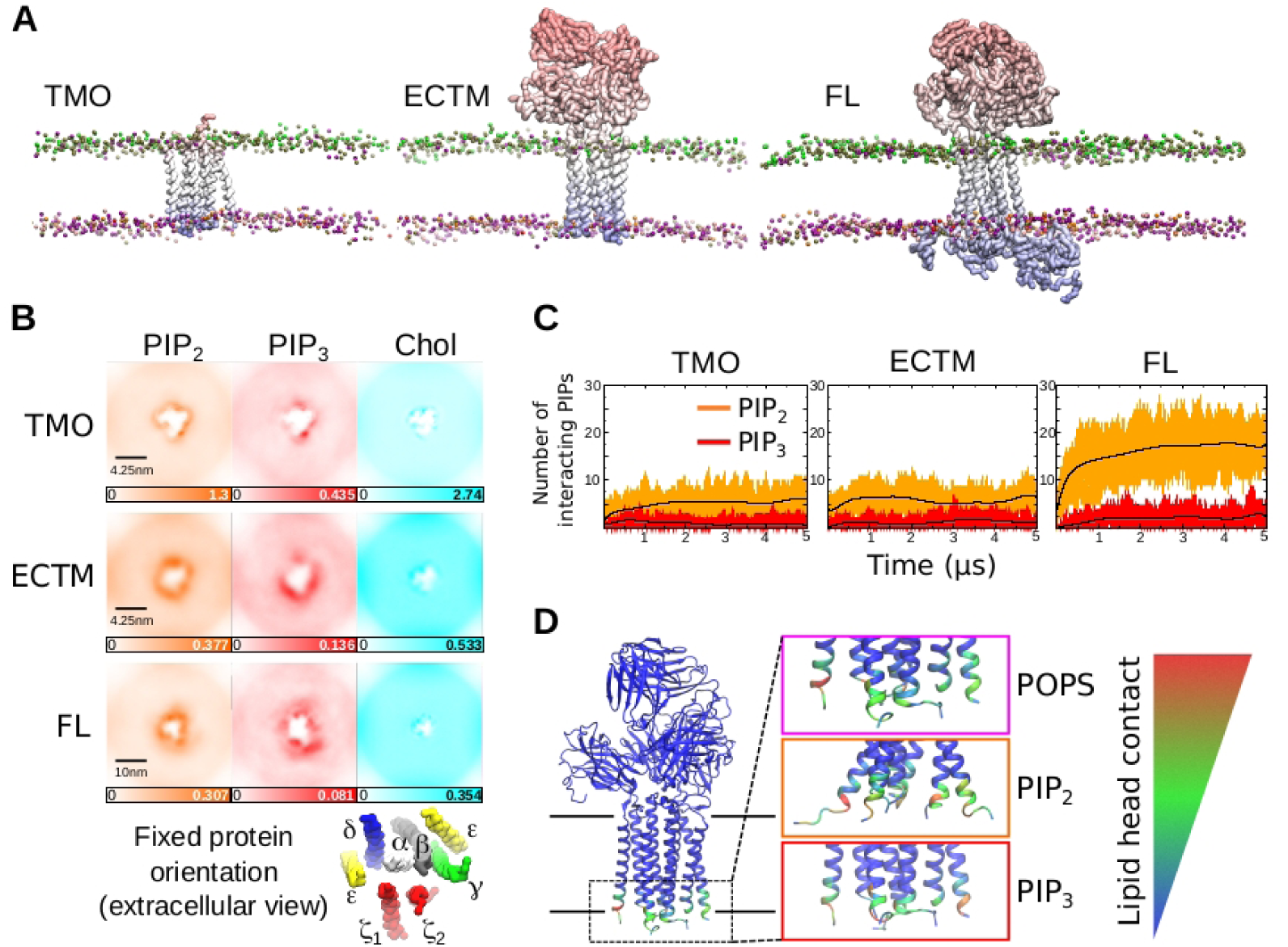
**(A)** Snapshots from the three sets of CGMD simulations (TMO, ECTM, FL) performed in this study. Here, only the protein and the lipid phosphate groups are shown for clarity. **(B)** Extracellular view of the average densities of PIP_2_, PIP_3_ and cholesterol molecules in the XY plane of the membrane from all five CGMD simulations combined. The protein was fixed in the center and its orientation is referenced below. **(C)** Number of interacting PIP_2_ or PIP_3_ lipids in the three CGMD systems (TMO, ECTM, FL) versus time. The smoothened black lines are a polynomial regression to the 10^th^ degree of the number of PIPs across time. **(D)** Interaction of the TCR’s cationic anchor with anionic lipids mapped onto the ECTM structure.

A distinct cholesterol annulus was clearly observed in the TMO simulations concentrated on the opposite side of the ζ ζ dimer, suggesting it to be a cholesterol-binding hotspot. Cholesterol bound to similar sites in the ECTM and FL simulations was also observed, but their density appears lower due to the concentration of unbound cholesterol in the membrane environment (**Fig 5B**). Nevertheless, we found that the average number of cholesterol molecules interacting with the TCR-CD3 were consistent in all three conditions (**Suppl Fig S4C**), suggesting that cholesterol binds to the TMRs independent of the ECDs or CYRs (**Suppl Fig S5**). Similarly, we quantified the change in the average number of PIP lipids contacting the TCR-CD3 across time. This revealed a three-fold increase in the number of PIP_2_ lipids in the FL simulations compared to the TMO and ECTM simulations (**Fig 5C**), suggesting that the CYRs played a significant role in enhancing PIP_2_ clustering. The number of interacting POPE lipids with the TCR-CD3 also increased in presence of the CYRs while the number of interacting POPS lipids showed a minor increase (**Suppl Fig S4C**). There were no differences in the average number of interacting PIP_3_ (**Fig 5B**), POPC and SM lipids (**Suppl Fig S4C**) when comparing the FL simulations to TMO and ECTM simulations.

We further questioned why there were some anionic lipid head-groups contacting the TCR-CD3 in the ECTM simulations despite the absence of the CYRs. This was due to the cationic residues present at the juxtamembrane region of TCR-CD3 (**Fig 5D**) (**Suppl Fig S6A**), collectively termed here as the ‘cationic anchor’. A multiple sequence alignment of the TMRs and the juxtamembrane region indicates that the cationic anchor is conserved across various species (**Suppl Fig S6B**). Consistently, our analysis suggested that the binding sites of PIP_2_, PIP_3_, and POPS (**Suppl Fig S7A, S7B, S8**) in the TMO and ECTM simulations occur largely in the juxtamembrane region of CD3ε and ζ subunits. In the FL simulations, binding of anionic head-groups occurred both in their juxtamembrane regions and CYRs. In comparison, the interaction of the TCR-CD3 with anionic lipid head-groups in the absence of its CYRs potentially suggest that the cationic anchor can alone retain one-third of anionic lipids around the TCR-CD3.

### Conformational changes and inter-chain interactions in the TCR-CD3

Inspection of the TMRs in the TMO, ECTM, and FL simulations at the end of 5 µs revealed a consistent loosening of the ζ 1 TMR from the CD3ε (δ) TMR (**Fig 6A, Suppl Fig S9A) (Suppl Movie M2**). Note that, in the cryo-EM structure (PDB:6JXR), their TMRs are in contact only at their N-terminal ends near the outer membrane leaflet surface. Calculation of the distance between the COM of ζ 1 and of CD3ε (δ) TMRs in the FL simulations showed that there was an increase in the distance between their TMRs (along the XY plane i.e. parallel to the membrane) compared to their initial distance calculated from the cryo-EM structure (**Suppl Fig S9B**). The untying of ζ 1 and CD3ε (δ) TMRs led to a decrease in distance between CD3δ and TCR β TMRs in all FL simulations (**Suppl Fig 9C**). During the simulations, the ζ 2 and CD3γ CYRs also formed contacts in addition to their TMR (**Suppl Fig S10A**). TCR αβ - ζ ζ interactions occurred mostly in the TMR with a very small number of interactions observed in the membrane-proximal extracellular region (**Suppl Fig S10B**).

**Figure 6.**
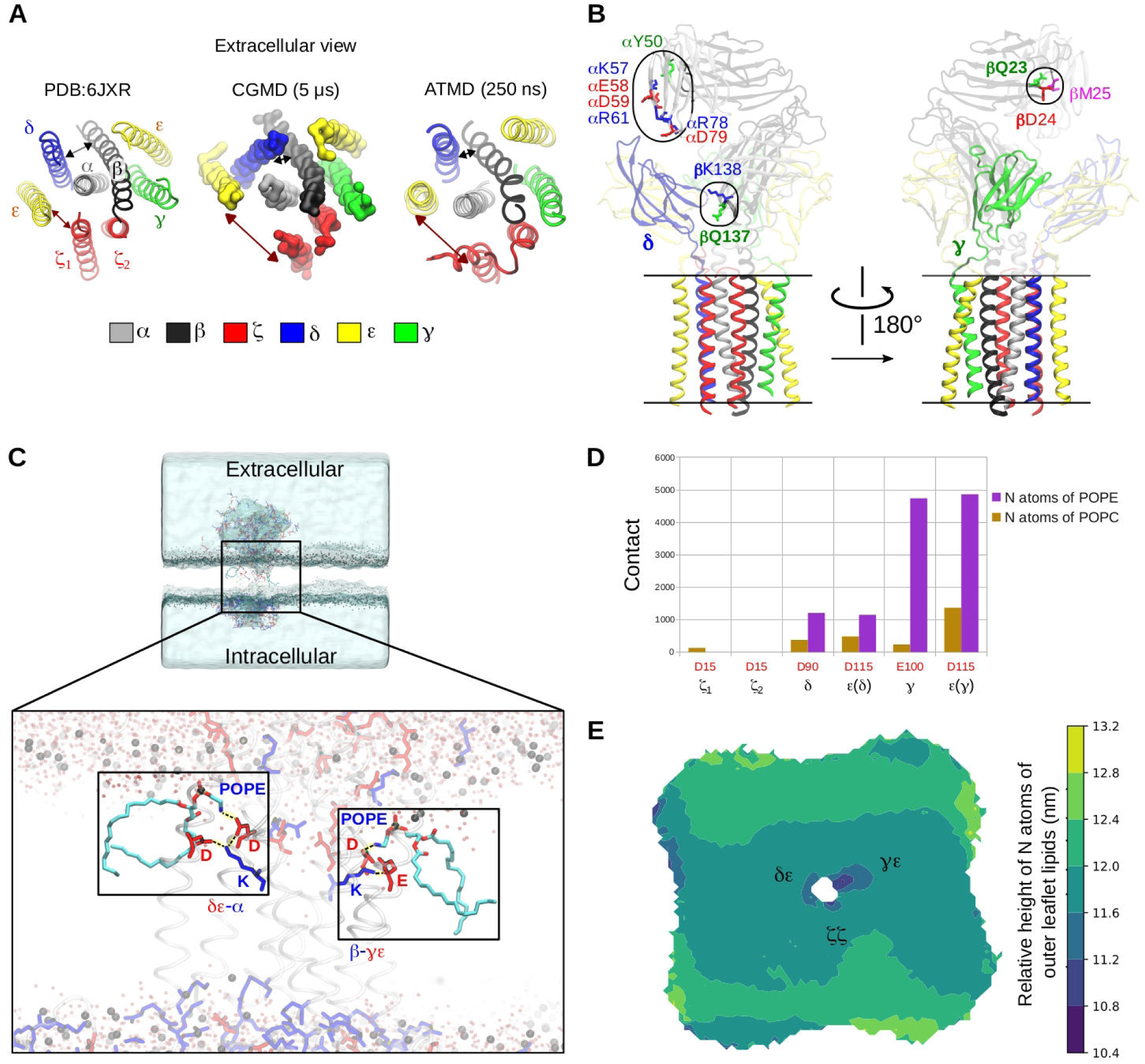
**(A)** Extracellular view of the TCR-CD3 TMR conformation from the cryo-EM structure (PDB:6JXR) compared to the most common conformation observed in CGMD simulations. ATMD simulations maintained the same conformation throughout the 250 ns simulation time. **(B)** New interactions of the TCR ; variable domain and of TCR β constant domain with CD3δ (left), and of the TCR β variable domain with CD3y seen in the ATMD simulations. **(C)** Snapshot from the end of one of our ATMD simulations of the entire TCR-CD3. The regions highlighted in boxes show pulled-in POPE head-groups whose amine (NH3) groups interact with the anionic sidechains of CD3 TMRs. The pulled-in POPE lipids are shown in cyan with their oxygen atoms shown in red and nitrogen atoms in blue. All hydrogen atoms are hidden for clarity. Anionic and cationic protein residues are represented as licorice sticks in red and blue, respectively. The TMR helices are shown as transparent cartoons. Water molecules are shown as transparent red spheres and the phosphorus atoms of all phospholipids are shown as black spheres defining the surface of the membrane. **(D)** Number of contacts of anionic residues of CD3 TMRs with nitrogen (N) atoms of POPC and POPE lipids from all ATMD simulations combined. **(E)** Extracellular view of the relative height of nitrogen atoms of all outer leaflet lipids (POPC, POPE, SM) mapped as a colour gradient from dark blue (low) to yellow (high). The TCR-CD3 TMR orientation is fixed in the center as shown in (A).

Although the cryo-EM structure shows that the CD3 ECDs are only contacted by the TCRa constant domains, our CGMD simulations revealed additional interactions of the TCRa variable domain with the CD3 ECDs (**Suppl Fig S10B**) indicating some flexibility within the TCR-CD3 ECDs (**Fig 2A top panel**). To analyse the flexibility of each TCR-CD3 ECD, we calculated the distances of the COM of (1) the αβ variable domain, (2) the αβ constant domain, (3) the δε, and (4) γ ε ECDs, each to a fixed point in the TMR (Cα atom of α R232 residue) along the vertical axis which is perpendicular to the membrane (**Suppl Fig S11A**). This showed that the TCR αβ variable domain (Vα Vβ) is the most flexible region of the TCR-CD3 ECD. This flexibility of Vα Vβ may be required for antigen recognition and initiating allosteric effects onto the rest of the complex.

### Atomistic molecular dynamics simulations (ATMD)

The CGMD simulations involved applying an elastic network (19) within each dimer (αβ, δε, γ ε, ζ ζ) to maintain their dimeric tertiary/quaternary structures during simulations, hence it restricted conformational changes within the dimers. Therefore, we also performed ATMD simulations of the entire TCR-CD3, thereby providing information on the protein-protein interactions and dynamics of the TCR-CD3 in atomistic detail (AT). From the end of the 5 µs CGMD simulations, the most common TCR-CD3 conformation, where ζ 1 and CD3ε (δ) TMRs had undergone loosening, was extracted along with the lipid environment and converted from CG to AT resolution (20). Three ATMD simulations were further performed for 250 ns each.

During the ATMD simulations, the TCR-CD3-lipid interactions observed in CGMD simulations were retained. The loosening between ζ 1 and CD3ε (δ) TMRs, seen in CGMD simulations was also maintained throughout the ATMD simulations suggesting that this conformation may be an energetically favourable configuration for the TCR-CD3. This loosening between ζ 1 and CD3ε (δ) TMRs brought CD3δ TMR closer to TCR β TMR as observed in CGMD simulations (**Fig 6A, Suppl Fig S9C**). Although ζ 1 and CD3ε (δ) TMRs made minimal contact, they interacted via their extracellular segments and CYRs. ζ 1 made no contact with the CD3γ ε dimer, while (2 interacted only with CD3γ via its TMR and CYR (**Suppl Fig S11B**). ATMD simulations revealed new interactions of Vα and Vβ with the ECDs of CD3δε and CD3γ ε respectively (**Fig 6B, Suppl Fig S11C**). In addition to the contacts revealed by the cryo-EM study (4), two more residues of TCRβ constant domain (β Q137, βK138) were found to interact with the CD3δ ECD (**Fig 6B, Suppl Fig S11C**). Furthermore, our ATMD simulations showed that the α-helical regions in the CYRs of ζ ζ were retained. Additionally, despite starting as unstructured regions, the CD3δ, ε, γ CYRs tended to form short α-helices in all ATMD simulations. CD3δ CYR had the smallest helical regions compared to all other CYRs (**Suppl Fig S12**).

ATMD of the TCR-CD3 complex in the membrane also revealed that, in the outer membrane leaflet, POPC and POPE head-groups were pulled toward the hydrophobic region of the membrane to ionically interact with anionic residues of the CD3 and ζ TMRs (**Fig 6C**). From all ATMD simulations combined, we calculated the number of contacts between the nitrogen atoms of all outer leaflet lipid head-groups i.e., POPC, POPE, SM, and the anionic residues of the CD3 and ζ TMRs. We found that the nitrogen atoms of SM made negligible number of contacts, POPC made some contacts, while POPE dominated interaction. The CD3γ ε TMR induced the most pulling of POPC/POPE head-groups into the membrane whereas ζ ζ TMR induced the least (**Fig 6D, 6E**). Moreover, the pulling down of lipid head-groups resulted in extracellular water solvating some charged residues of the TCR-CD3 TMR, thereby leading to a local membrane deformation in the outer membrane leaflet. Previous MD studies proposed that this occurs prior to TCR-CD3 assembly (21) but our studies suggest that this occurrence is also a feature of the entire TCR-CD3 complex.

## Discussion

The CYRs of the CD3 and ζ subunits are essential to mediate T cell activation, as pMHC binding induces tyrosine phosphorylation of their ITAMs by LCK and intracellular signal propagation by the ZAP-70 tyrosine kinase (7). Biochemical and biophysical studies have suggested that the CYRs of the CD3ε and ζ subunits of a non-stimulated TCR-CD3 complex interact with the inner leaflet of the plasma membrane (9, 10, 22). These data may imply that TCR-CD3 ligation induces dissociation of the CYRs from the inner leaflet of the membrane (11, 23). This dissociation allows augmented ITAM access by active LCK (13) followed by sustained ITAM phosphorylation leading to activation of ZAP-70 to propagate intracellular signalling (7). Therefore, it is possible that ligand-induced conformational changes in the TCR-CD3 facilitate ITAM exposure. It has also been proposed that the dissociation of CYRs could occur by ITAM phosphorylation due to an increase in their net electronegative charge. However, the clusters of positively charged amino acids strongly interacting with the membrane may compete with this dissociation mechanism (8).

Our study shows that the CYRs of CD3δε, CD3γ ε and ζ ζ dimers exhibit a coiled and interlaced conformation forming contacts with each other. This coiled conformation of the tails allowed only some cytoplasmic tyrosines to penetrate into the membrane whilst the sidechains of some other cytoplasmic tyrosine were hidden; their sidechains pointed inwards into the coiled conformation of the CYRs. The transient exposure of these hidden ITAM tyrosines to intracellular solvent during our simulations supports findings that resting T-cells can undergo basal phosphorylation and signalling (24, 25). Moreover, our simulations suggest that the network of protein-protein interactions among the CYRs reduce the solvent accessibility surface area, thereby reducing the probability of ITAM phosphorylation in its resting state. Therefore, an allostery-based stimulation of conformational changes in the TCR-CD3 which alters the interactions of ζ ζ TMR with the rest of the TCR-CD3 as seen in our simulations can potentially contribute to the unbinding of ζ ζ CYRs from the CD3 CYRs and further increase the exposure of their ITAMs for phosphorylation. Our data also shows that the overall SASA of tyrosine sidechains on each ζ CYR is larger than that of CD3 CYRs. This is in agreement with a previous study that suggested that ζ ζ ITAM multiplicity can enhance signalling (26).

Recent cryo-EM studies revealed most of the TCR-CD3 quaternary structure in a detergent environment, i.e. its TMRs and ECDs, but not its CYRs (4). This allowed us to employ molecular modelling to complete the TCR-CD3 structure and simulate its dynamics in a bilayer that closely mimics the lipid composition of a TCR-CD3 activation domain in its native plasma membrane (14). We observed in our simulations that the TCR-CD3 conformation was somewhat divergent from the cryo-EM structure, indicating that its conformation may differ in a membrane environment compared to that obtained in a detergent environment. Our simulations suggested an alteration in the TMR configuration where the ζ ζ and CD3δε dimers lost contact thereby bringing CD3δ and TCR β TMRs closer than observed in the cryo-EM structure. Moreover, we identified interactions of Vα Vβ with CD3 ECDs indicating flexibility of the antigen-binding domain of the TCR αβ, in line with studies that identified allosteric sites in the TCR αβ constant domains (2, 27-30).

This work also demonstrates that the TCR-CD3 creates a unique lipid fingerprint in the membrane by forming selective interactions with anionic head-groups of PIP lipids. This is in agreement with experimental findings that have suggested that PIP lipids regulate membrane dynamics and TCR-CD3 activation (31). Such unique membrane footprints have also been suggested for other membrane proteins (32). Our simulations show that PIP lipids interact strongly with the BRS of CD3ε and ζ subunits in the intracellular region, consistent with findings suggesting that BRS mediate interaction with PIPs and modulate signalling (9, 10, 22). Moreover, the BRS of CD3ε is suggested to serve as a docking site for LCK (33, 34) whereas that of ζ is reported to induce membrane-binding of its tails (12, 22) and help localise TCRs at the immunological synapse (10). Similar cationic patches were observed in the juxtamembrane regions of receptor kinases (35) and other signalling receptors such as Epha2 (36), EGF receptors (37), and integrin-talin complexes (38, 39) where they were shown to play critical functions in receptor activation by interacting with anionic lipids.

In this study, we also show that the cationic residues situated at the interface of the TMR and CYR of the TCR-CD3 can alone maintain an anionic lipid environment suggesting that the TCR-CD3 complex can maintain an anionic environment, albeit smaller, even when its cytoplasmic tails are dissociated from the membrane. The presence of the positively charged cytoplasmic tails enhances the formation of a distinct annulus of PIP lipids around the TCR-CD3. The triggering of PIP clustering may further create a suitable lipid environment for recruiting peripheral proteins such as LCK. Mutation studies also showed that the LCK-SH2 domain interacts with PIP_2_ lipids via a cationic patch at K182 and R184, which is distinct from its phospho-tyrosine binding site (40). The SH2 domain of ZAP-70, which is homologous to LCK-SH2, was also found to bind to PIP_2_ head-groups and phospho-tyrosines independently. It was also found that these SH2 domains cannot associate with stimulated TCRs when their lipid-binding site is mutated (41) suggesting that lipid interaction of SH2 domains could be critical to their association with the TCR-CD3. Therefore, the formation of an anionic lipid environment enriched in PIP_2_ around the TCR-CD3 shown in our study may be key for its interactions with LCK and other SH2 domain-containing protein kinases.

In summary, we propose the first molecular model of the entire resting TCR-CD3 complex inserted in an asymmetric bilayer containing lipids found in the TCR-CD3 activation domain. Our dynamic model suggests selective interactions of the TCR-CD3 with anionic head-groups, especially those of PIP_2_, whose interactions were enhanced in the presence of CD3 and (CYRs. The clustering of PIP_2_ lipids around TCR-CD3 may facilitate the initial interaction of its CYRs with LCK, participate in TCR-CD3 clustering and aid in the spatial organization of the immunological synapse. Therefore, our findings can lead to further studies, e.g. involving mutations in the CYRs to alter protein-lipid interactions, to better understand the molecular mechanism of TCR activation and signalling.

## Methods

### Molecular Modelling

The cryo-EM structure (PDB:6JXR) (4) was used as a template to obtain the complete TCR-CD3 model. Sequences of each subunit were obtained from UniProtKB (uniprot.org): ζ:P20963, δ:P04234, ε:P07766, γ:P09693, α:A0A0B4J271, β:P0DSE2. Modeller 9.2 was used to model the entire TCR-CD3 complex (42, 43) (**Suppl Fig S1**) along with UCSF Chimera (44). Hydrogen atoms were added to the model and topologies were generated using Gromacs 2016 (45).

### Coarse-grained molecular dynamics (CGMD) simulations

The structural models were coarse-grained using the martinize script. CGMD simulations were conducted using Gromacs version 5.0 with the Martini forcefield (46, 47). The TCR-CD3 is comprised of the αβ, δε, γ ε, ζ ζ dimers non-covalently bonded with each other. To replicate this, an elastic network model (19) with a cut-off distance of 0.7 nm was applied within each dimer to maintain their dimeric tertiary/quaternary structures. The protein complex was then placed in a simulation box and inserted into a complex asymmetric bilayer using the *Insane* tool (48). The concentrations of lipids in both bilayers (**table 1**) are based on lipidomics studies of TCR-CD3 activation domains (14). CG waters were added and the system was neutralised with 0.15M of NaCl. They were energy minimised using the steepest descent algorithm followed by equilibration with the protein backbone particles position-restrained for 2 ns. The equilibrated system was used to generate systems with differing initial velocities for five production simulations run for 5 µs each with 20 fs time-step. Every frame in each simulation was generated at 200 ps intervals. The barostat and thermostat used for CGMD production simulations were Parinello-Rahman (1 bar) (49) and V-rescale (50), respectively. A compressibility of 3×10^−4^ bar^-1^ was used. The LINCS algorithm (51) constrained bond lengths.

### Atomistic molecular dynamics (ATMD) simulations

A commonly observed structural conformation of the entire TCR-CD3 along with its lipid environment was extracted from the end of a CGMD simulation and backmapped (20) to AT resolution. Gromacs 2016 with the Charmm36 forcefield (52) was used for backmapping. Before performing ATMD, the simulation box size was reduced along the vertical (Z) axis to prevent simulating excess solvent particles, thus minimizing the computational cost. The protein-lipid conformation was retained during the resizing of Z axis of the simulation box. The TIP3P water model was used along with 0.15M of NaCl to neutralize the system. The system was energy-minimised with the steepest descent algorithm followed by a step-wise (0.25 → 1 → 1.5 → 1.8 → 1.9 → 2 fs time-step) NPT equilibration with the protein backbone position-restrained. The final equilibration step was conducted for 2 ns with a 2 fs time-step. The equilibrated system was then used to generate systems with different initial velocities for three repeat production simulations each of which were run for 250 ns with a 2 fs time-step. Every frame in each simulation was generated at 40 ps intervals. The V-rescale thermostat (323K) and Parrinello-Rahman semi-isotropic barostat (1 bar) (49) was used with a compressibility of 4.5×10^−5^ bar^-1^. The LINCS algorithm (51) applied constraints on bond lengths and the Particle Mesh Ewald algorithm (53) defined long-range electrostatics. A summary of simulations is shown in **table 2**.

**Table 2.** Summary of simulations conducted in this study.

| Simulations | Resolution | Particles | Simulation box<br>(X × Y × Z axis) | Duration | Replicas |
| --- | --- | --- | --- | --- | --- |
| TCR TMO + complex bilayer | CG | 26728 | 25 × 25 × 21 nm | 5 μs | 5 |
| TCR ECTM + complex bilayer | CG | 54106 | 25 × 25 × 21 nm | 5 μs | 5 |
| TCR FL + complex bilayer | CG | 196303 | 25 × 25 × 21 nm | 5 μs | 5 |
| TCR FL + complex bilayer | AT | 1145375 | 25 × 25 × 21 nm | 250 ns | 3 |

### Analysis

Calculation of all protein-lipid, protein-protein and protein-solvent contacts in the CGMD simulations used a 0.55 nm distance cut-off to define a contact. The same cut-off value was used to calculate contacts of protein residues with hydrophobic lipid acyl chains. Similarly, all contact analyses for ATMD simulations used a 0.4 nm cut-off. All interaction profiles represent merged data from all simulation repeats and were performed using *gmx mindist* command. Solvent accessibility of intracellular tyrosines was calculated using *gmx sasa* considering 10 ns intervals starting from 1 µs time from all CGMD simulations combined. To calculate clusters of TCR cytoplasmic conformations, and lipid densities around the protein, the trajectories of all simulation repeats were concatenated using *gmx trjcat* and the protein orientation was fixed in the center using *gmx trjconv*. The *gmx densmap* and *gmx xpm2ps* commands were used to produce lipid density images. The clustering analysis of the cytoplasmic conformations was performed using the *gmx cluster* command. Clustering analysis was done using a 0.35 nm cut-off for the protein backbone RMSD for every 10^th^ frame i.e. 400 ps, resulting in 10,000 structures grouped into 389 clusters. Distance and radius of gyration analyses were performed using the *gmx distance* and *gmx gyrate* commands respectively. VMD was used for visualisation and rendering snapshots of the molecular system. The RMSD trajectory tool of VMD (54) was also used to perform alignments of TMR helices. The electrostatic potential of the entire TCR-CD3 was obtained using the APBS electrostatics tool (55) integrated with VMD. The MUSCLE tool was used to perform multiple sequence alignments (56). Calculation of the relative height of lipid nitrogen atoms was based on their positions along the vertical (Z) axis i.e. perpendicular to the membrane. For this calculation, the TCR orientation was fixed in the center in each ATMD simulation before concatenating all of them. The python script to perform this calculation was obtained from https://github.com/jiehanchong/membrane-depth-analysis. The *gmx do_dssp* command was used to calculate the secondary structure formations in the TCR-CD3 CYRs considering 1 ns intervals from all ATMD simulations combined.

## Acknowledgments

A.C.K. was supported by a Springboard Award from the Academy of Medical Sciences (Great Britain) and the Wellcome Trust (grant number: SBF002\1031). O.A. was supported by a Wellcome Trust grant (WT200844/Z/16/Z). This research was enabled in part by support provided by ARC3 and ARC4 supercomputers, part of the High Performance Computing facility at the University of Leeds, Leeds, United Kingdom.

## Supplementary Figures

**Figure S1.**
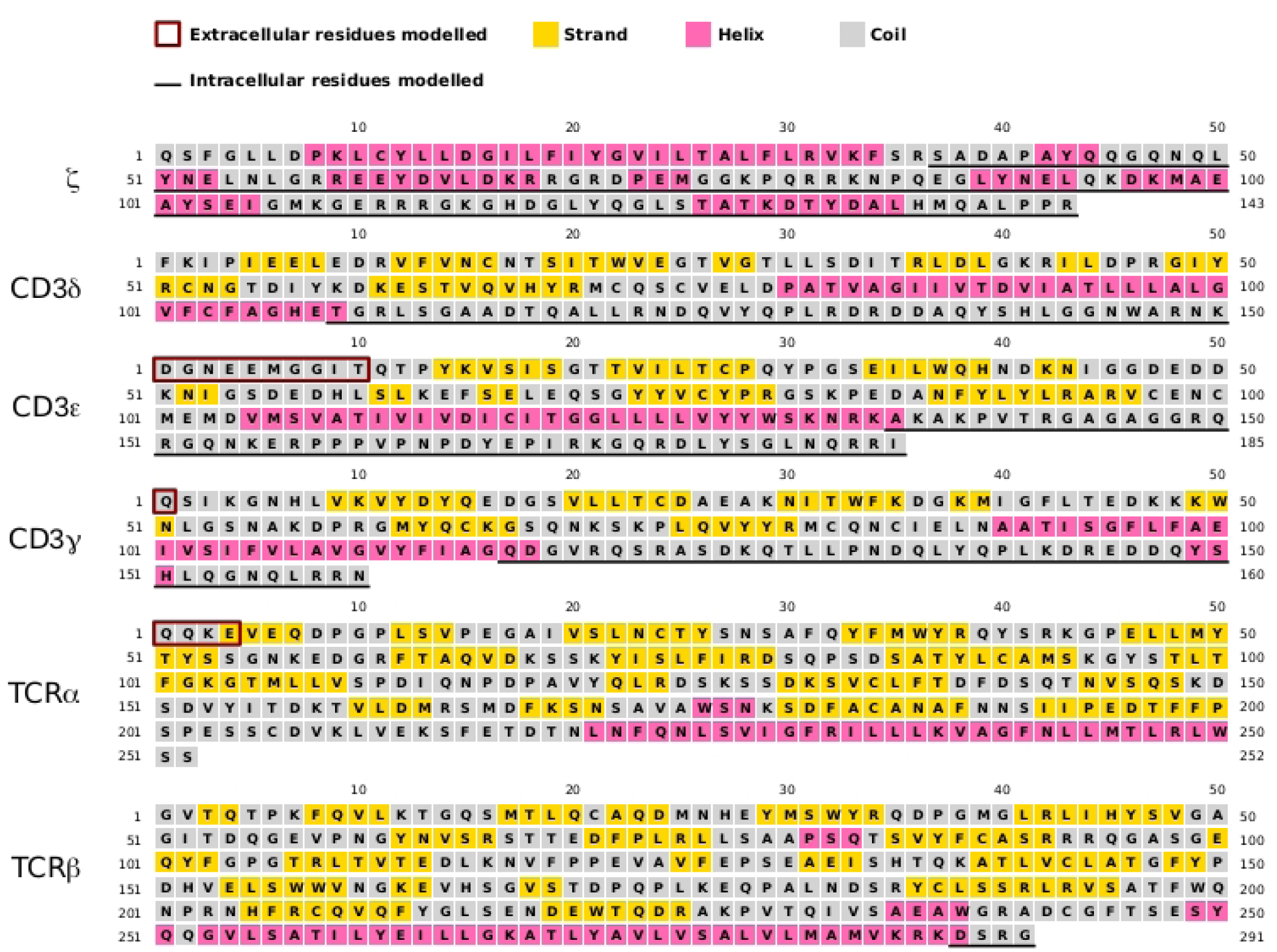
Sequences of each full-length subunit of the TCR-CD3 and their secondary structure predictions as suggested by the PSIPRED 4.0 server. The extracellular and intracellular residues modelled in this study are shown in boxes and underlined respectively.

**Figure S2.**
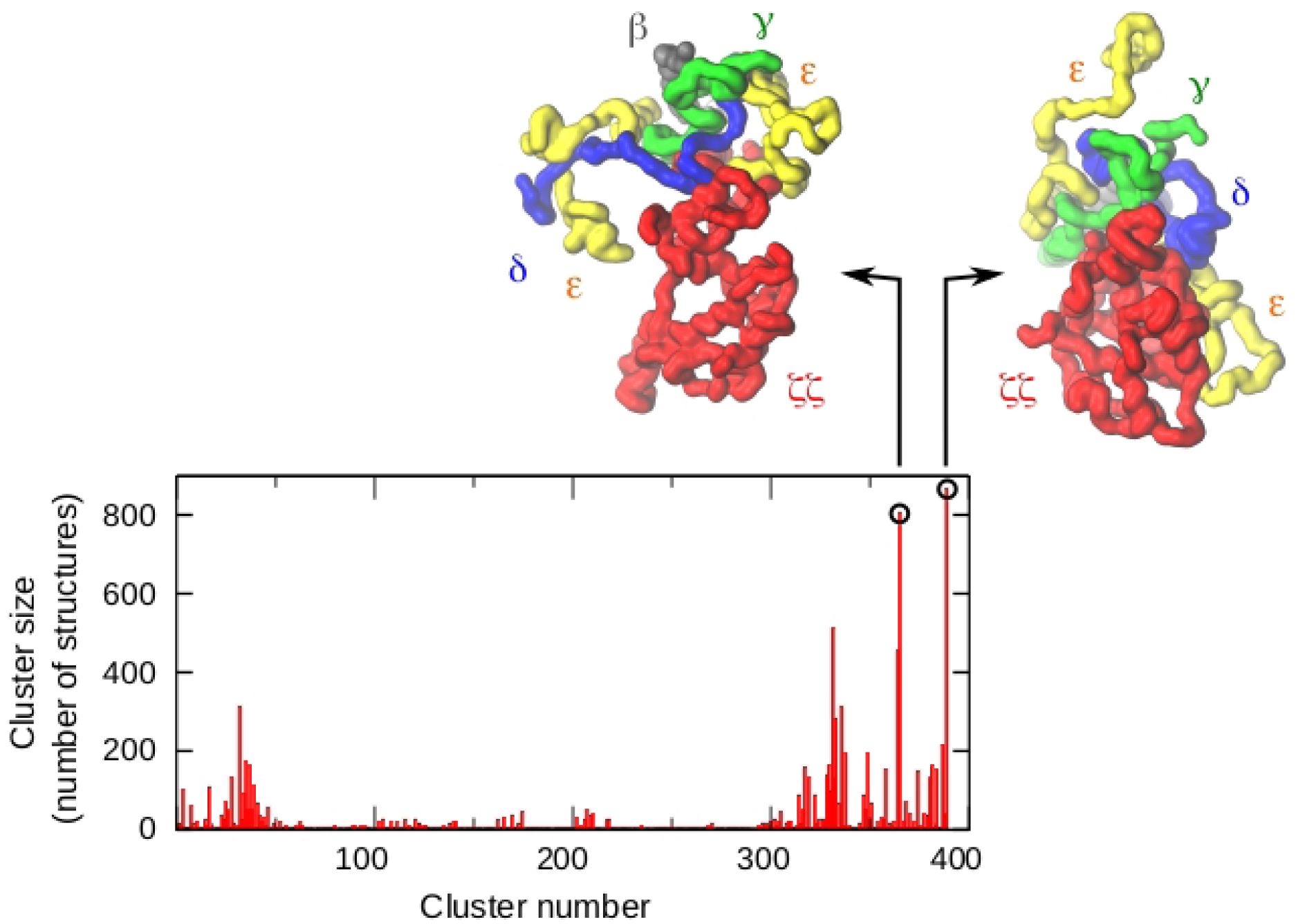
TCR-CD3 cytoplasmic conformations grouped into clusters based on a 3.5Å RMSD cut-off. The clusters containing the highest number of structures indicate the most stable conformation of the TCR-CD3 cytoplasmic region. The structure is coloured by subunit: ζ:red, δ:blue, ε:yellow, γ:green, α:silver, β:black.

**Figure S3.**
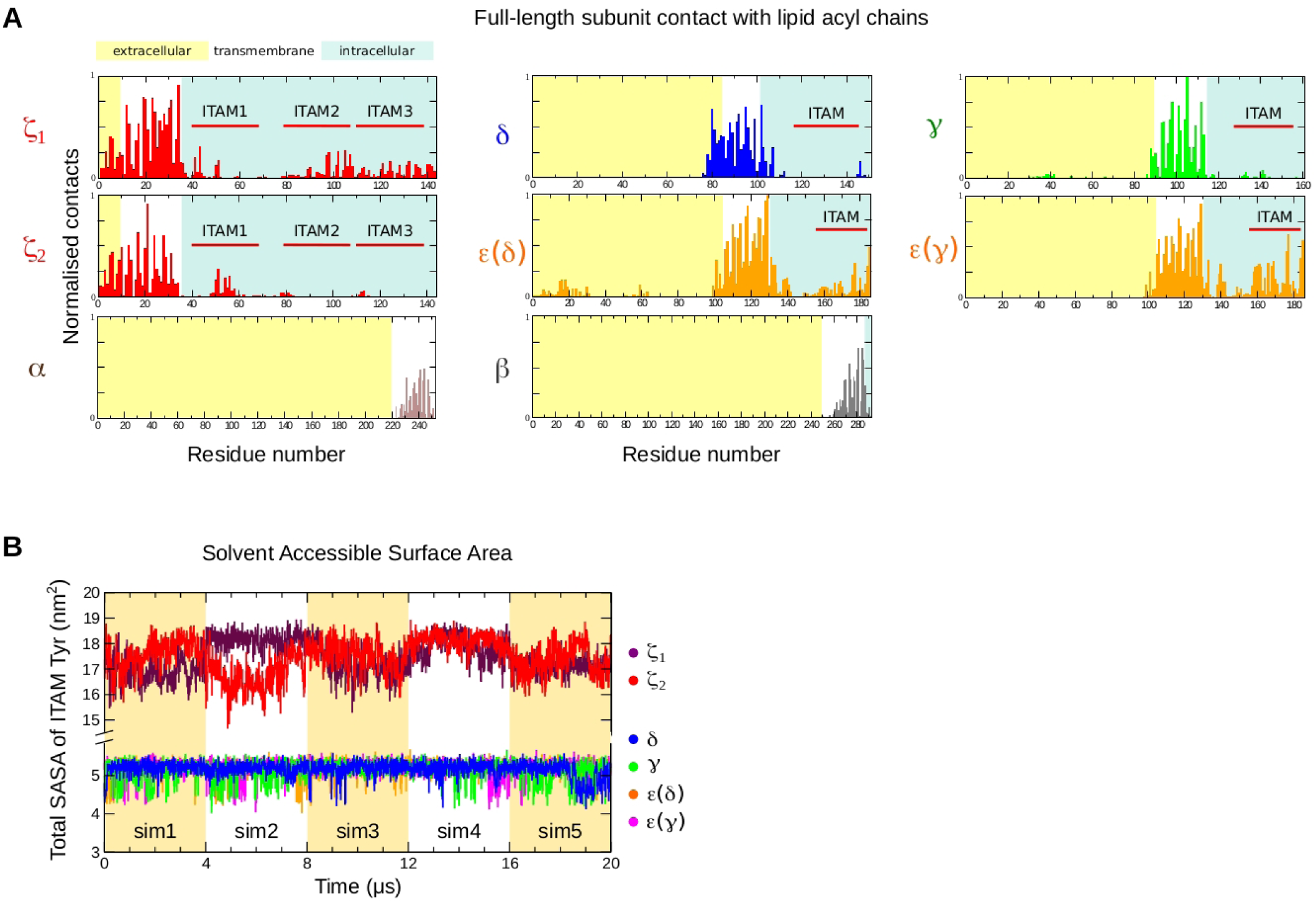
**(A)** Normalised number of contacts of each full-length TCR-CD3 subunit with hydrophobic lipid acyl chains, from all five CGMD simulations. Contacts are normalised by dividing the number of contacts of each residue by the highest number of contacts across all subunits. Therefore, the value 1 represents the highest contact while O represents no contact. **(B)** The solvent accessibility surface area (SASA) of all lTAM tyrosines combined from each CD3 and ζ subunit.

**Figure S4.**
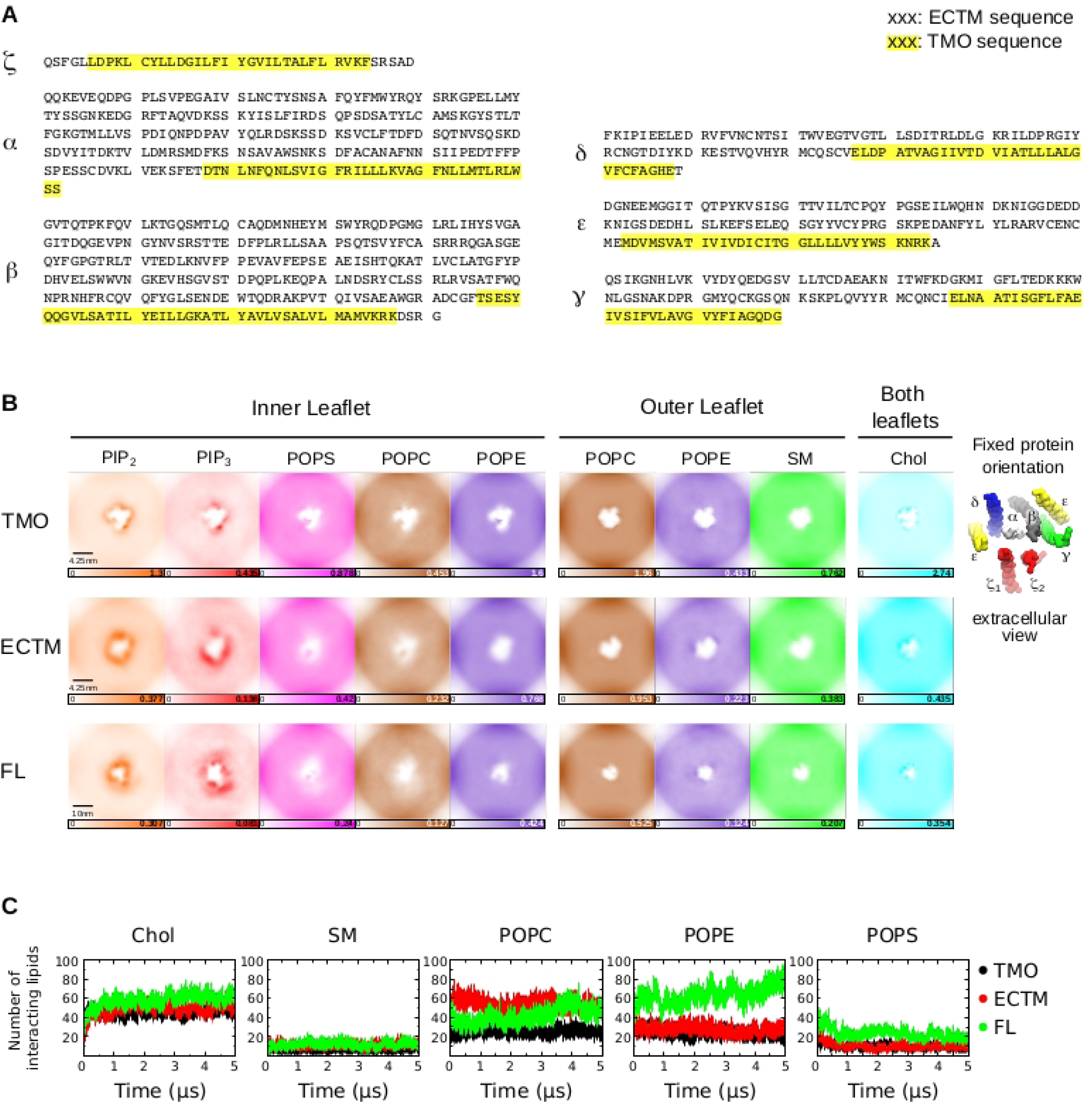
**(A)** Sequences of the TCR-CD3 used for CGMD simulations of the ECTM system (ectodomain and transmembrane only), and of the TMO system (transmembrane only). **(B)** Extracellular view of the density of each lipid type in the membrane when the orientation of the TCR-CD3 is fixed in the center. The colour gradient scale for each density displays the number of lipids corresponding to the minimum and maximum value. **(C)** The average number of cholesterol, SM, POPC, POPE, and POPS lipids interacting with the TCR-CD3 over 5 µs time from all CGMD simulations ofTMO, ECTM, FL systems.

**Figure S5.**
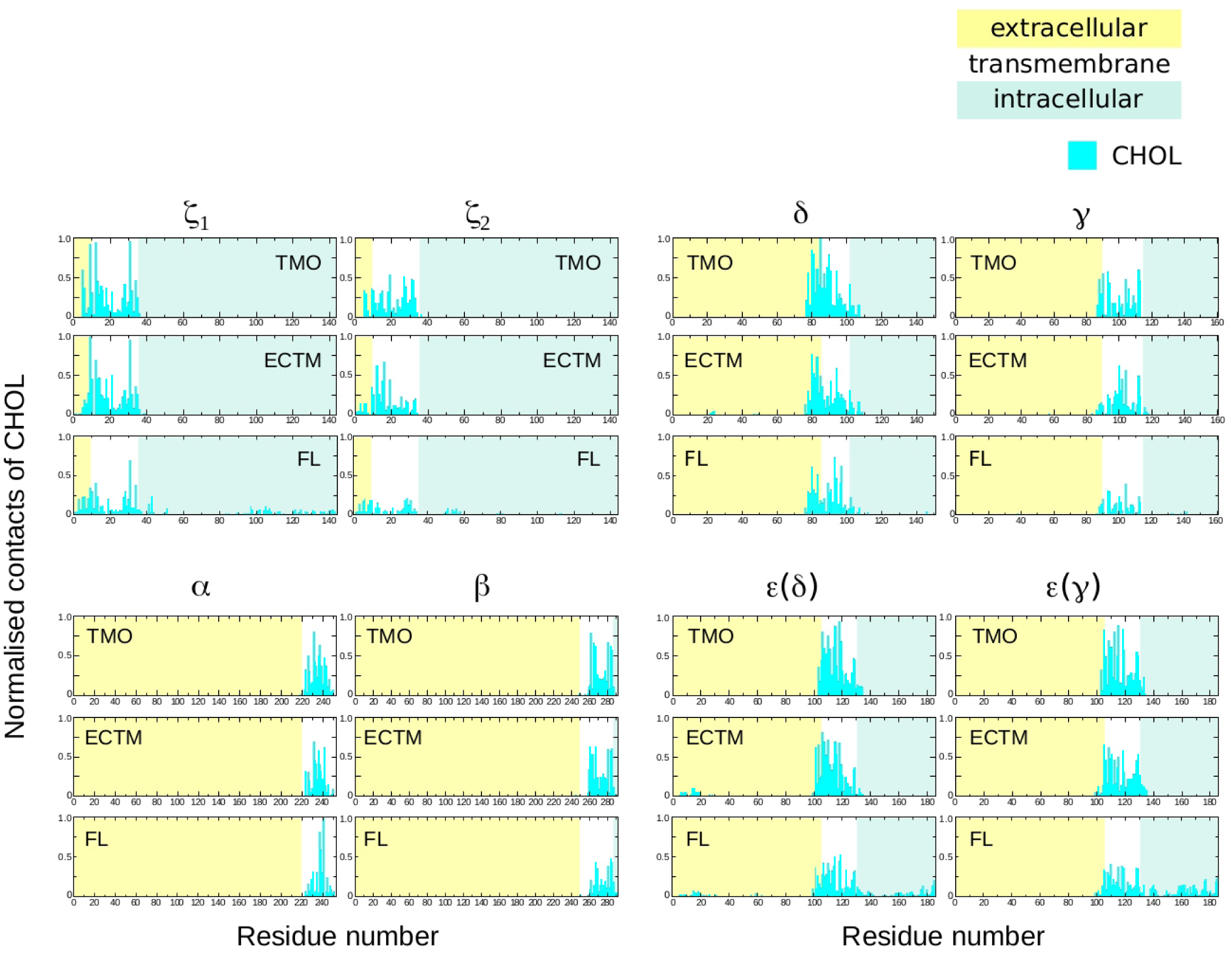
Normalised contacts of all TCR-CD3 subunits with cholesterol. Normalisation is done by dividing the contacts of each residue in the TCR-CD3 by the highest number of contacts. Therefore, the value 1 represents the highest contact while 0 represents no contact. The three TCR-CD3 systems: TMO, ECTM, FL, are normalised separately.

**Figure S6.**
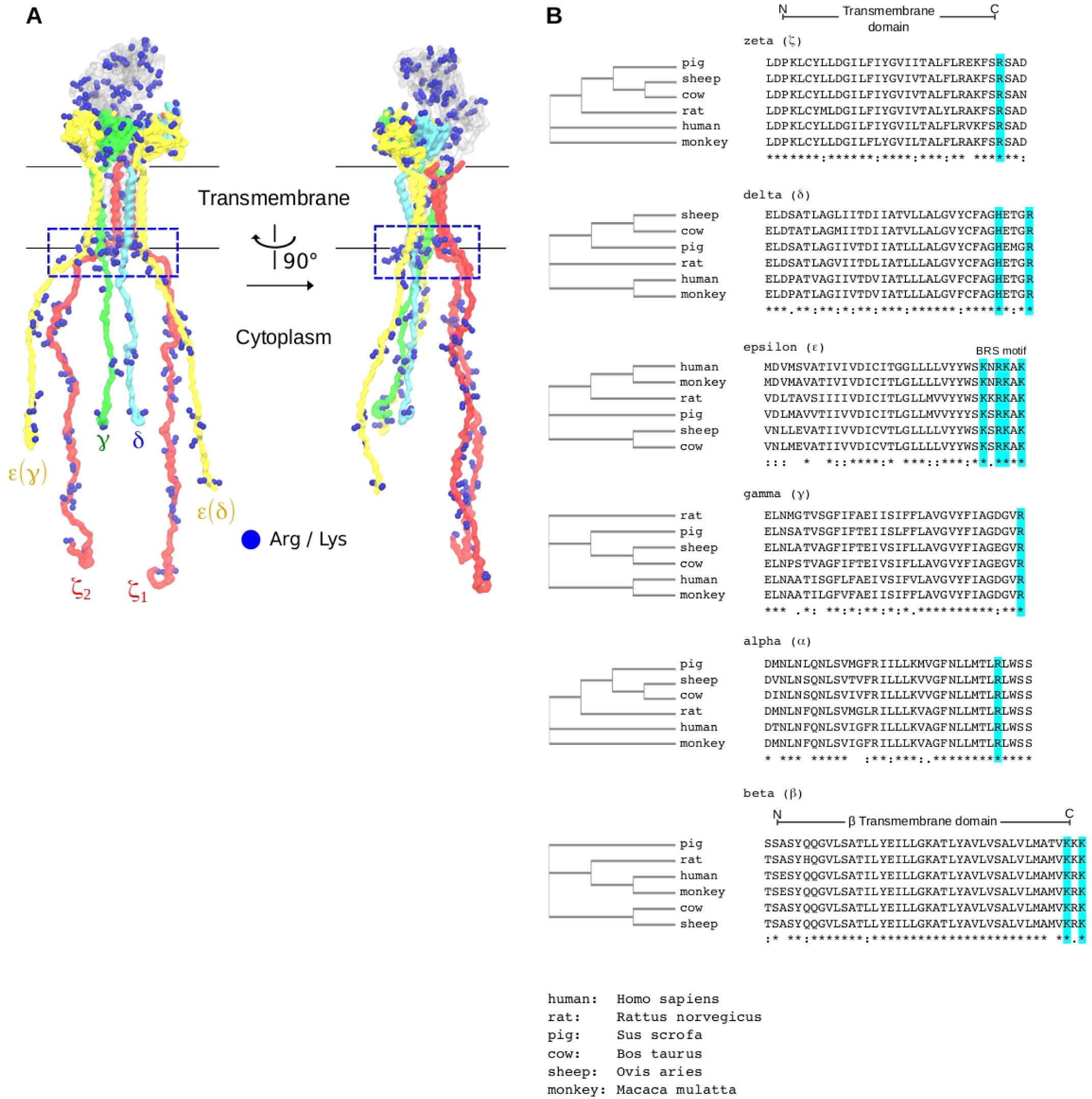
**(A)** The cationic anchor of the TCR-CD3 at the interface of the TMR and cytoplasmic region is indicated within a box. **(B)** A multiple sequence alignment of the TMR and juxtamembrane residues of all TCR-CD3 subunits indicating the conservation of cationic residues at the TMR-cytoplasmic interface across different species.

**Figure S7.**
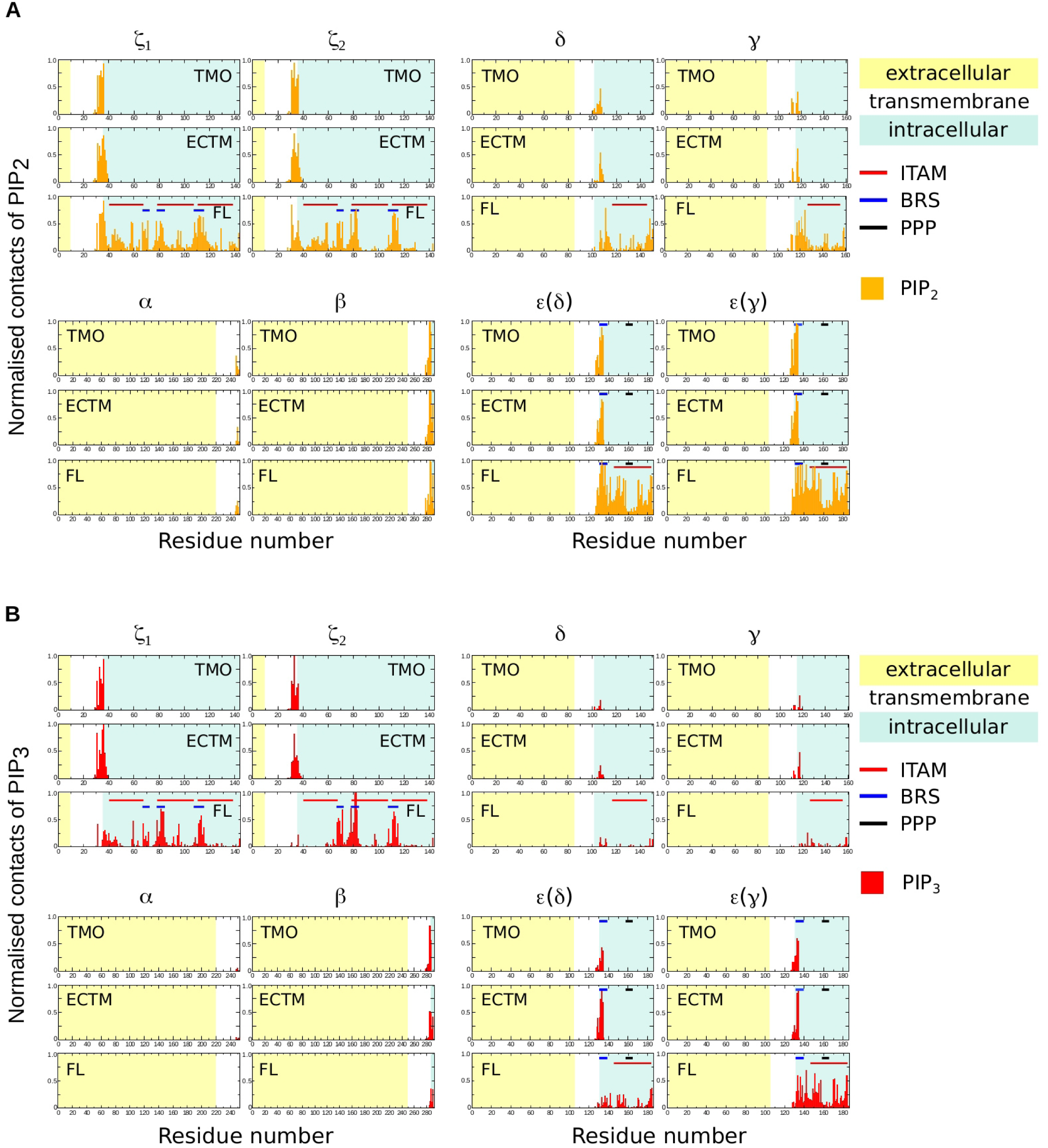
Normalised contacts of all TCR-CD3 subunits with **(A)** PIP_2_ and **(B)** PIP_3_. Normalisation is done by dividing the contacts of each residue in the TCR-CD3 with a lipid type by the highest number of contacts with that lipid type. Therefore, the value 1 represents the highest contact while 0 represents no contact. The three TCR-CD3 systems: TMO, ECTM, FL, are normalised separately.

**Figure S8.**
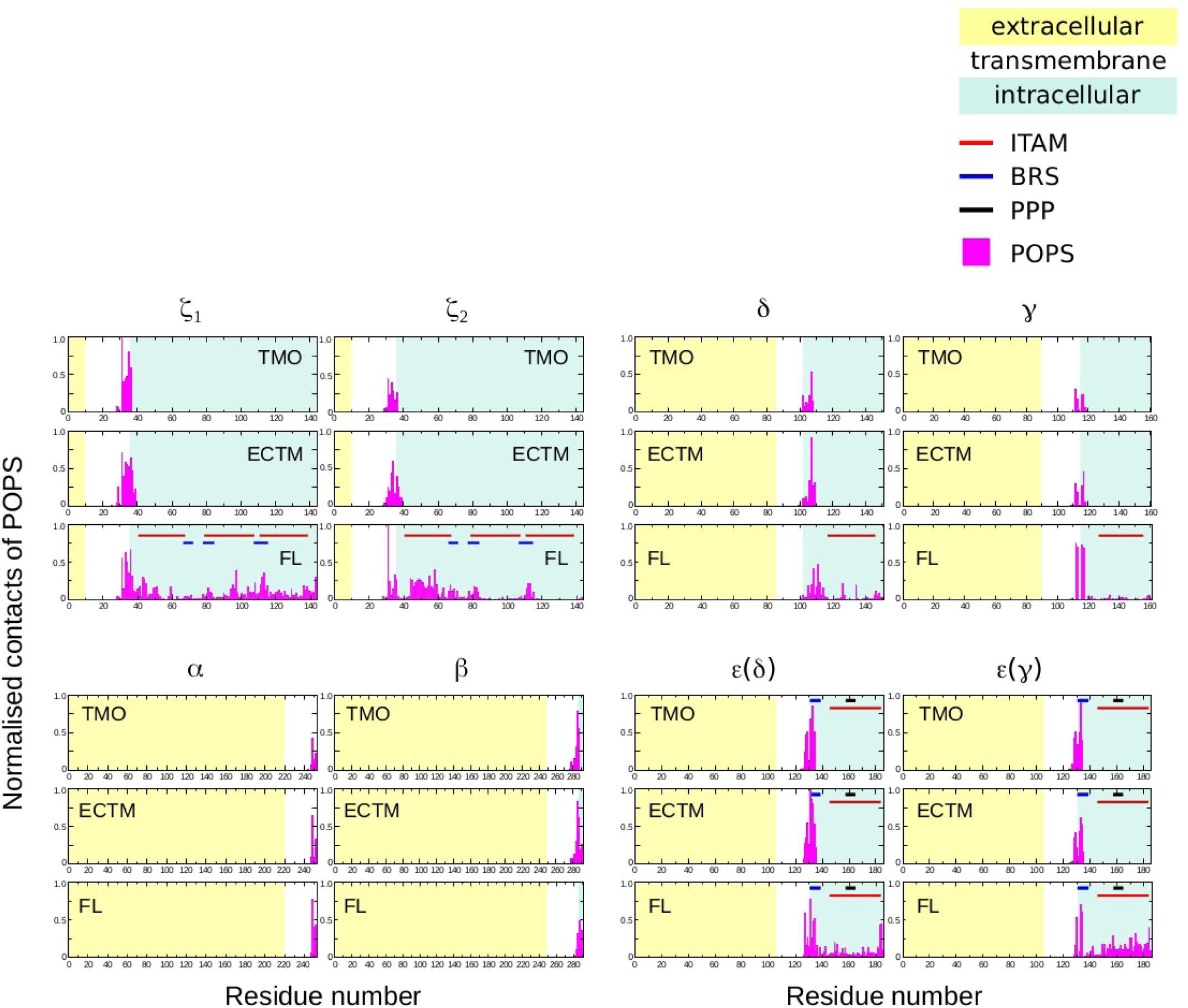
Normalised contacts of all TCR-CD3 subunits with POPS. Normalisation is done by dividing the contacts of each residue in the TCR-CD3 with POPS by the highest number of contacts with POPS. Therefore, the value 1 represents the highest contact while 1 represents no contact. The three TCR-CD3 systems: TMO, ECTM, FL, are normalised separately.

**Figure S9.**
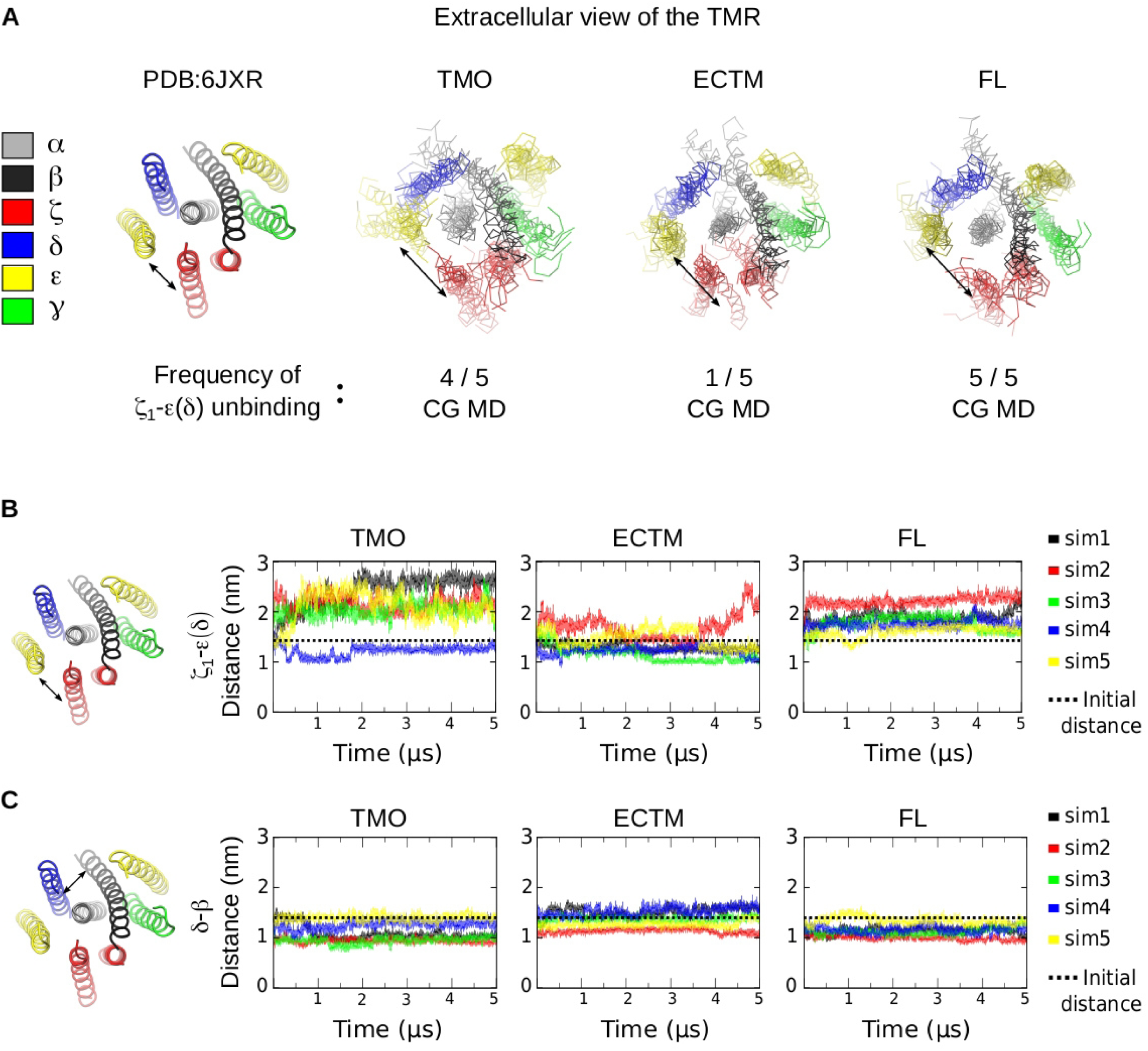
**(A)** Extracellular view of five TMR snapshots aligned from the end of the TMO, ECTM, FL simulation systems indicating the frequency of ζ 1-ε (δ) dissociation compared to the cryo-EM TMR structure (PDB:6JXR). **(B)** Distance between the center of mass of ζ 1 and of ε (δ) subunits, and **(C)** distance between the center of mass of δ and of β subunits in all simulations systems over 5 µs compared to their initial distances calculated from the cryo-EM structure.

**Figure S10.**
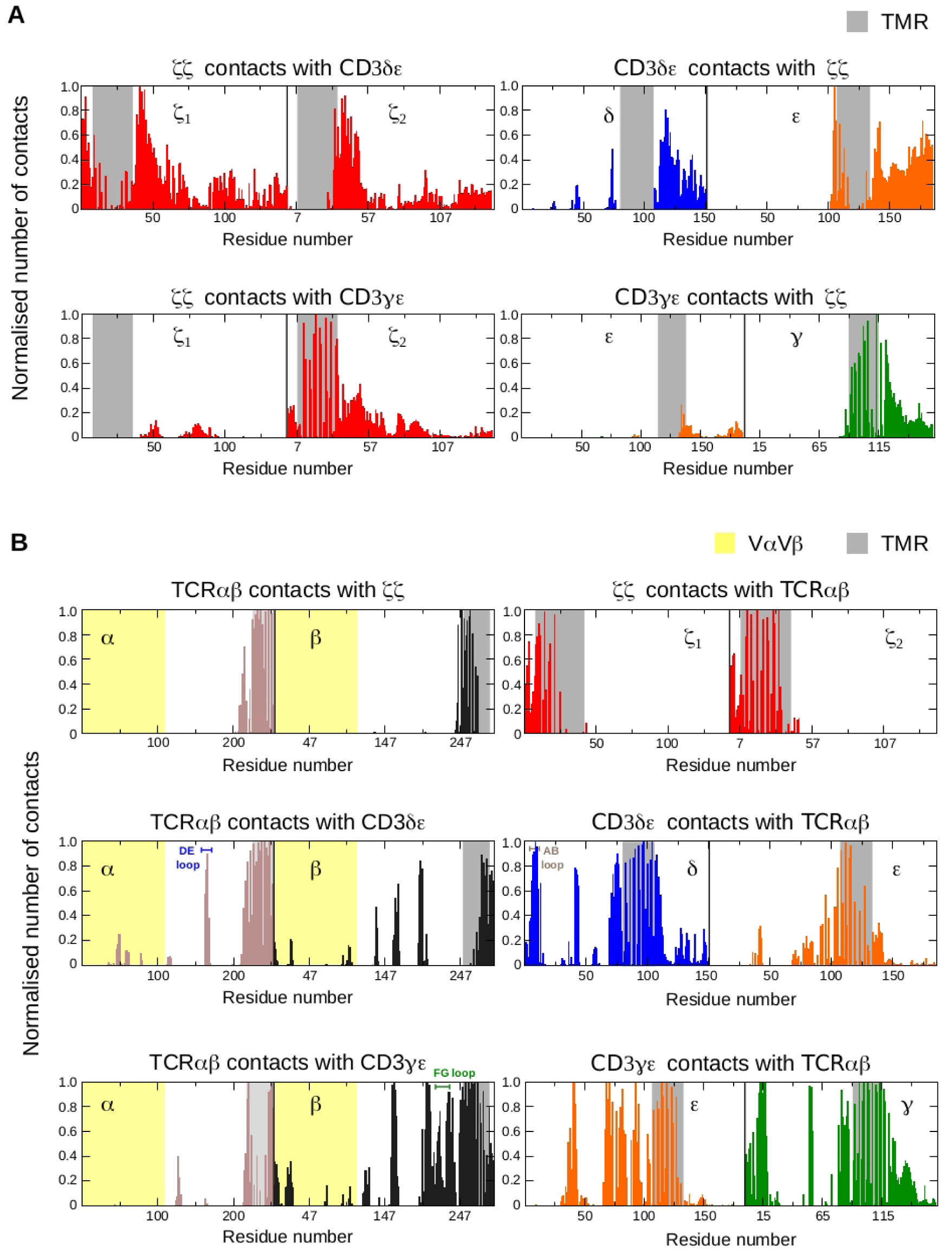
Normalised full-length inter-chain interactions of **(A)** the ζ ζ dimer with the CD36 δε and CD3γ ε dimers, and **(B)** the TCR αβ dimer with the ζ ζ, CD3γ ε, and CD3γ ε dimers in CGMD simulations. Normalisationis done by dividing the contacts of each residue by the highest number of contacts within each dimer.

**Figure S11.**
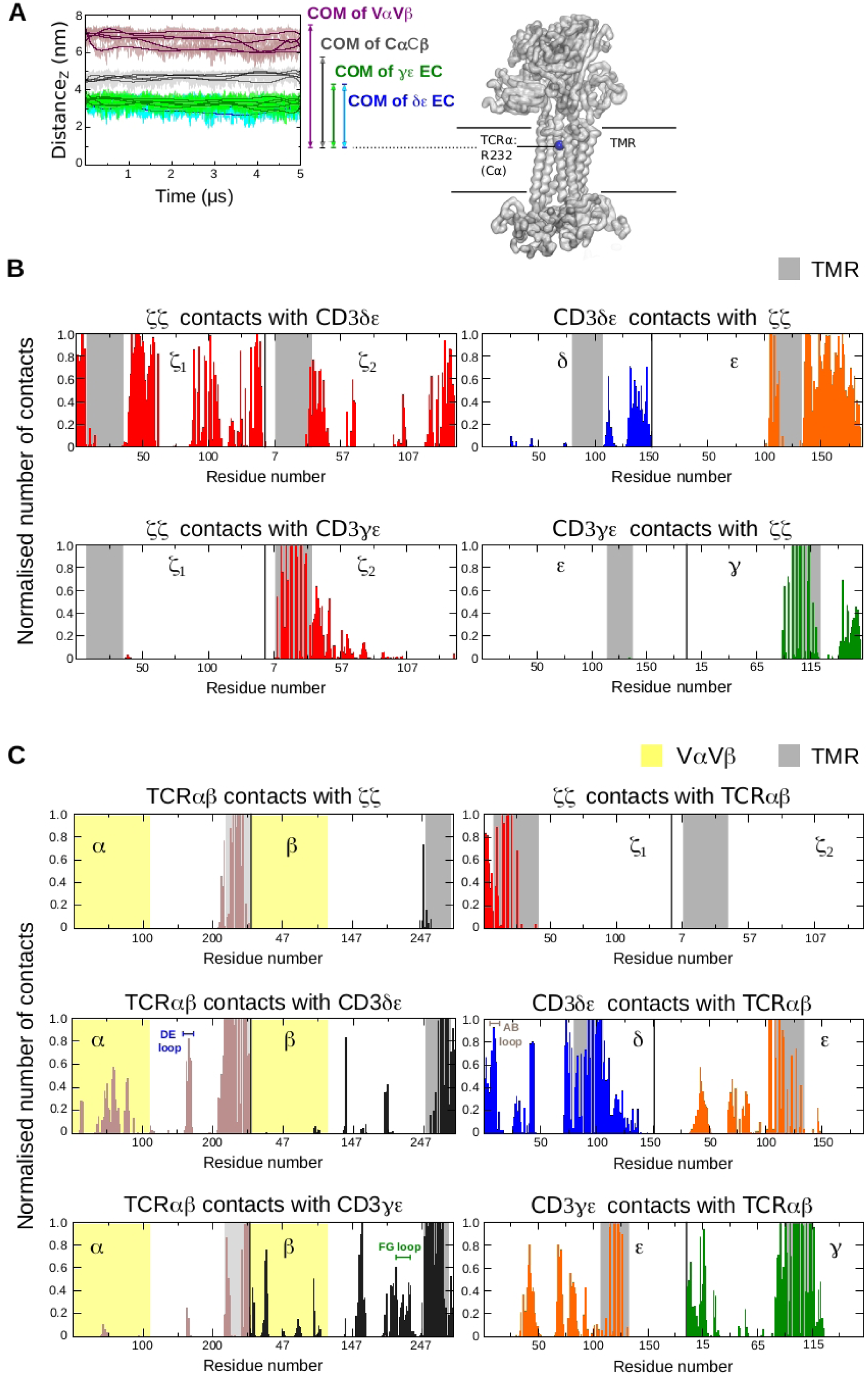
**(A)** Distance of the center of masses of the TCR αβ) variable domain (Vα Vβ), the TCR αβ) constant domain (Cα Cβ), the CD3δε, and CD3ζ ε ectodomains (EC), each to the Cα atom of αR232 residue in the TMR along the vertical (Z) axis. **(B)** Normalised full-length inter chain interactions of the ζ ζ dimer with the CD3δε and CD3ζ ε di mers, and **(C)** the TCR αβ) dimer with the ζ ζ, CD3δε, and CD3ζ ε dimers in ATMD simulations. Normalisation is done by dividing the contacts of each residue by the highest number of contacts within each dimer.

**Figure S12.**
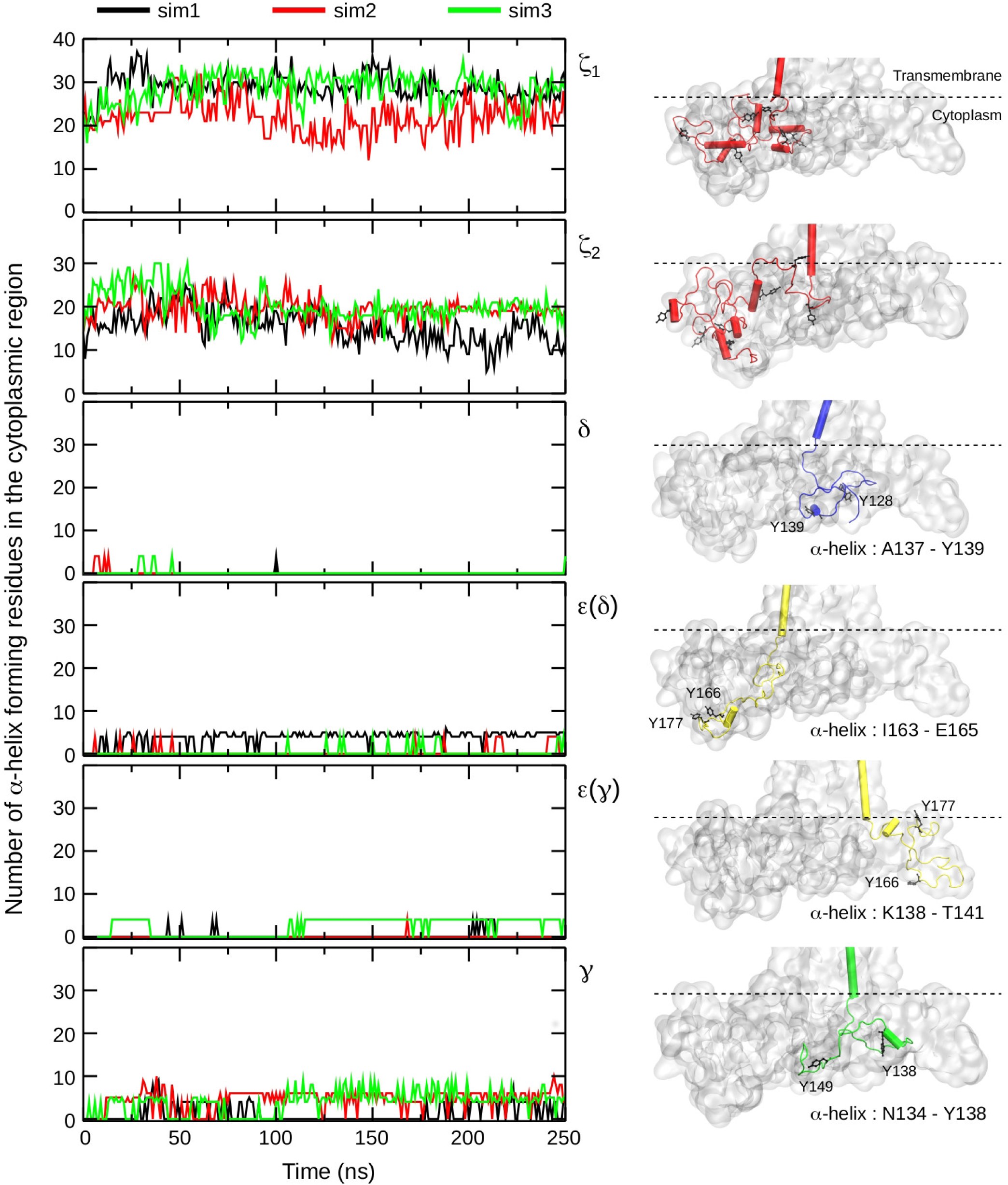
Number of α -helix forming residues in the cytoplasmic region of CD3 and ζ subunits versus time from all three ATMD simulations (left). Location of cytoplasmiCα -helices relative to the ITAM tyrosines of the respective subunits (right). The ITAM tyrosines and α -helix forming residues observed at the end of 250 ns from simulation-3 are labelled. The entire protein is shown using the surface representation, the ITAM tyrosines are represented as black ball and sticks, and the subunits are shown in cartoon representation and coloured as follows: ζ red, δ : blue, ε yellow, γ green (right).

